# Inflammatory IL-1 signaling remodels epidermal stem cell compartments by suppressing Wnt activity

**DOI:** 10.64898/2026.02.06.704488

**Authors:** Hung Manh Phung, Ikuto Nishikawa, Nguyen Thi Kim Nguyen, Aiya K. Yesbolatova, Ahmed M. Hegazy, Tomson Kosasih, Jun Aoi, Satoshi Fukushima, Sho Hiroyasu, Hitoshi Takizawa, Aiko Sada

**Affiliations:** Laboratory of Skin Regeneration and Aging, International Research Center for Medical Sciences (IRCMS), Kumamoto University, Kumamoto, Japan; Division of Skin Regeneration and Aging, Medical Institute of Bioregulation, Kyushu University, Fukuoka, Japan; Zoology Department, Faculty of Science, Minia University, El-Minia, Egypt; Laboratory of Stem Cell Stress, International Research Center for Medical Sciences (IRCMS), Kumamoto University, Kumamoto, Japan; Department of Dermatology and Plastic Surgery, Faculty of Life Sciences, Kumamoto University, Kumamoto, Japan; Department of Dermatology, Graduate School of Medicine, Osaka Metropolitan University, Osaka, Japan

**Keywords:** Epidermal stem cells, Slow-cycling cells, Inflammation, Stem cell heterogeneity, IL-1 signaling, Canonical Wnt pathway

## Abstract

The skin epidermis is maintained by spatially organized stem cell populations with distinct cellular dynamics; however, how inflammation affects this heterogeneity remains largely unknown. Here, we demonstrate that acute skin inflammation alters epidermal stem cell compartments through IL-1-mediated suppression of canonical Wnt signaling. Lineage tracing in inflamed mouse skin revealed that slow-cycling Dlx1^+^ epidermal stem cell clones persist, whereas fast-cycling Slc1a3^+^ clones decline through enhanced differentiation and lineage conversion, driving the reorganization of epidermal stem cell compartments. IL-1 signaling is both necessary and sufficient for this change: administration of IL-1α/β recapitulates these effects, while transgenic induction of the IL-1 decoy receptor preserves the balance of stem cell populations. IL-1 suppresses canonical Wnt activity in both the mouse epidermis and human keratinocytes, and Wnt ligand administration restores the fast-cycling compartment *in vivo*. Together, these results identify a reversible IL-1–Wnt axis that governs inflammation-induced stem cell plasticity and spatial tissue remodeling.

**Highlight:**

- Inflammation induces reversible remodeling of epidermal stem cell compartments
- Distinct epidermal stem cell populations exhibit differential responses to inflammation
- IL-1 suppresses canonical Wnt signaling, thereby biasing fast-cycling stem cell behavior
- Reactivation of Wnt signaling restores stem cell population balance under inflammatory conditions

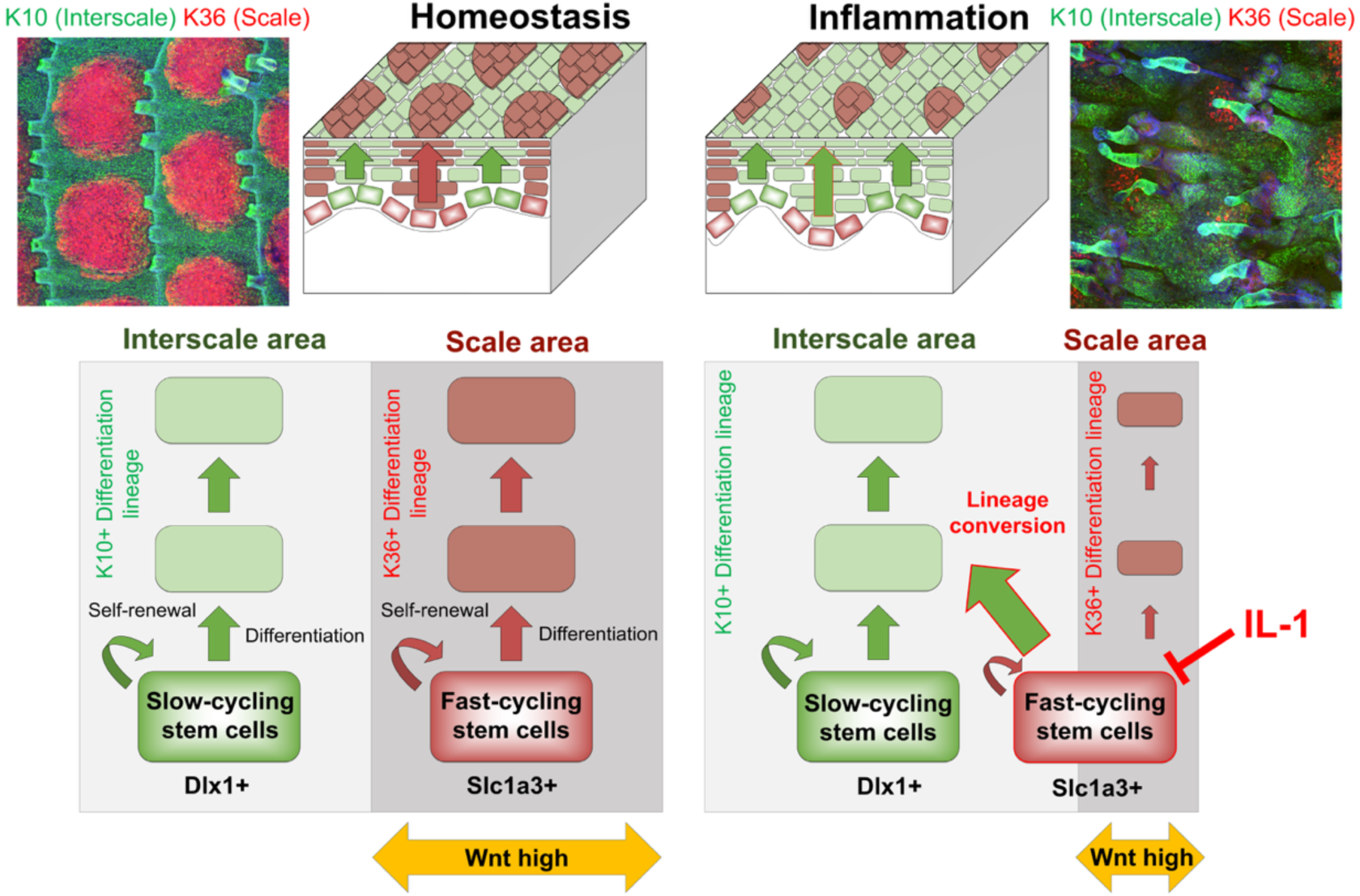

## Introduction

Adult tissues are maintained by spatially organized populations of tissue-resident stem cells that exhibit distinct molecular signatures^1,2^. This stem cell heterogeneity is regulated by intracellular signaling and stem cell–intrinsic transcriptional/epigenetic programs, enabling distinct stem cell populations to adapt to physiological demands, including tissue damage, environmental stress, and aging^1,2,3,4,5,6^. Among these extrinsic cues, inflammatory stress has emerged as a potent modulator of stem cell behavior across different tissues^7,8,9,10^. Experience of inflammation in the skin can be retained as transcriptional and epigenetic memory in some contexts, altering subsequent responses to injury and stress^11,12,13,14^. However, how inflammatory stress affects coexisting heterogeneous stem cell populations and the mechanisms underlying these cell-type-specific responses remain poorly understood.

The mouse tail epidermis provides a unique model for dissecting tissue stem cell dynamics under inflammatory stress. This tissue comprises two anatomically and molecularly distinct regions: the interscale, maintained by Dlx1^+^ slow-cycling epidermal stem cell population, and the scale, maintained by Slc1a3^+^ fast-cycling epidermal stem cell population^15,16,17,18^. Clonal lineage tracing and transcriptomic studies have identified other stem/progenitor populations, indicating a broader heterogeneity within the tail epidermis^19,20,21,22,23,24,25,26,27,28^. Notably, this heterogeneity is not randomly distributed but spatially organized into anatomically defined compartments, in which distinct stem cell populations generate characteristic lineage outputs^15,16^. These compartmentalized stem cell populations show differential responses to tissue damage, external stimuli, oncogenes, and aging^16,29,30,31,32,33,34^. Thus, this spatial compartmentalization provides a tractable framework for interrogating how inflammatory stress differentially impacts coexisting stem cell populations and addressing their contribution to epidermal homeostasis and remodeling. Importantly, analogous compartmentalization and stem cell heterogeneity are observed in the human skin epidermis and oral mucosa, where inter-ridge and rete ridge structures parallel the mouse interscale and scale^35,36,37^, highlighting the broader relevance of this model.

Proinflammatory cytokines, such as interleukin-1 (IL-1), TNF, and interferons, act as both modulators of immune responses and regulators of tissue stem/progenitor cell behavior^38,39^. These signals can reprogram the cellular homeostatic state, leading to clonal bias, impaired self-renewal/differentiation, or exhaustion of tissue stem cells, as previously demonstrated in hematopoietic and intestinal stem cells^8,10,38^. IL-1 is a key driver of inflammatory responses to infection and tissue damage^40^. IL-1 signaling is mediated by IL-1α and IL-1β, which bind IL-1R1 and activate signaling through the adaptor protein MyD88^41^. This pathway is counterbalanced by IL-1R2, a decoy receptor that sequesters IL-1 ligands and limits excessive IL-1 responses^42^. In the skin, IL-1 plays diverse roles: cooperating with EGFR, thereby driving keratinocyte proliferation^43^; enhancing epidermal γδ T-cell expansion, triggering hair follicle stem cell proliferation^44^; promoting adipocyte progenitor activation during wound repair^45^; and stimulating regenerative responses via IL-1R/Myd88 signaling^46^. However, how inflammatory stress reshapes the balance, fate, and spatial organization of heterogeneous epidermal stem cell populations remains largely unknown.

In this study, we addressed how IL-1-dependent skin inflammation alters the composition and behavior of epidermal stem cell populations. By combining lineage tracing, transcriptomic, and functional analyses with genetic models, we demonstrated that IL-1 signaling suppresses canonical Wnt activity under inflammatory conditions, preferentially compromising the maintenance of the Slc1a3⁺ fast-cycling population while preserving the Dlx1⁺ slow-cycling population. This change is reversible and can be attenuated by blocking IL-1 signaling or reactivating Wnt signaling. Together, these findings demonstrate that inflammatory IL-1 signaling transiently overrides homeostatic Wnt control, thereby selectively reshaping the balance and spatial organization of heterogeneous epidermal stem cell populations.

## Results

### Inflammatory stress induces reversible reorganization of epidermal stem cell compartments

To examine how inflammation influences epidermal stem cell populations, we employed an established mouse model of acute skin inflammation via the topical application of 12-O-tetradecanoylphorbol-13-acetate (TPA) to the tail skin (Fig. 1A, B). TPA triggers a robust inflammatory response that mimics features of various skin pathologies, including epidermal hyperplasia, leukocyte infiltration, and increased proinflammatory cytokine (IL-1, IL-6, and TNF-α) production through activation of protein kinase C (PKC)-dependent cascades in epidermal keratinocytes (Fig. 1C)^47,48,49,50,51,52^. Consistently, we observed increased cytokine expression (Fig. 1D), epidermal thickening (Fig. 1E, G), and increased Ki-67^+^ proliferative cells in the epidermis (Fig. 1F, H) at days 3 and 9 after the initiation of TPA treatment.

**Figure 1.**
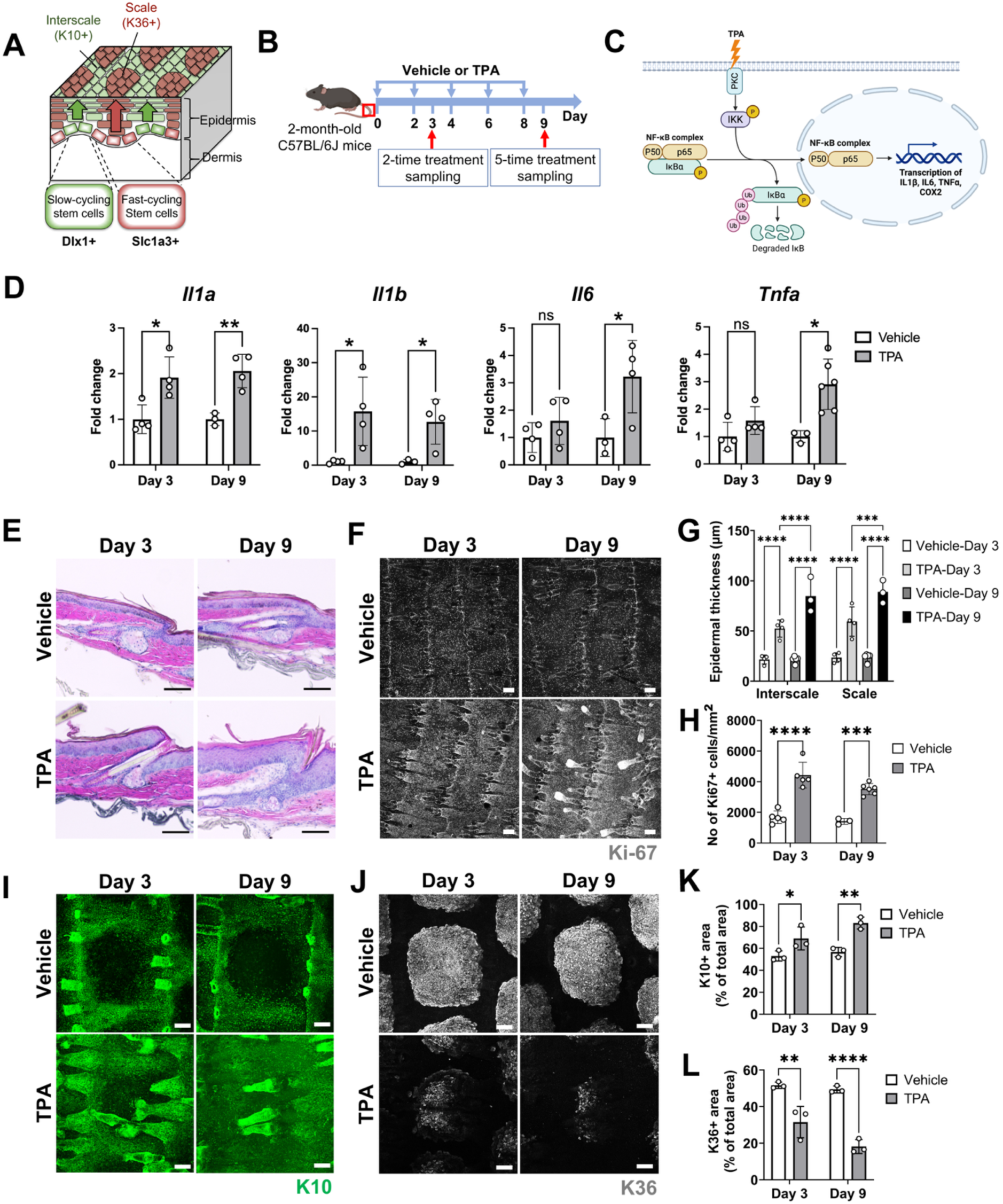
Inflammatory stress remodels epidermal stem cell spatial compartments. (**A**) Schematic representation of the interfollicular epidermis of mouse tail skin. Dlx1^+^ slow-cycling epidermal stem cells produce the K10^+^ interscale lineage (green), whereas Slc1a3^+^ fast-cycling epidermal stem cells produce the K36^+^ scale lineage (red). (**B**) Experimental scheme of 12-O-tetradecanoyl-phorbol-13-acetate (TPA)-induced skin inflammation. Ethanol (vehicle) or TPA was applied topically to the mouse tail skin every other day, and the skin was dissected on day 3 or day 9. (**C**) Molecular pathway of TPA-induced skin inflammation. (**D**) Expression of inflammatory cytokines in the whole tail skin by qRT–PCR. Expression levels were normalized to the corresponding vehicle-treated controls at each time point (day 3 or day 9). (**E, G**) Hematoxylin and eosin staining of sagittal sections of tail skin (E). Scale bars: 100 µm. Epidermal thickness was measured in the interscale and scale regions (G). (**F, H**) Whole-mount immunostaining of Ki-67 (F) and its quantification (H). Scale bars: 100 µm. (**I–L**) Whole-mount staining of K10 (interscale, I) and K36 (scale, J), and their quantifications (K, L). Scale bars: 100 µm. All data are presented as the mean ± SD. Each dot represents an independent biological replicate. Statistical significance was assessed using one-way ANOVA for (D) and two-way ANOVA for (G, H, K, L). *, *p* < 0.05; **, *p* < 0.01; ***, *p* < 0.001; ****, *p* < 0.0001; ns, not significant.

We next assessed how this inflammatory state impacts the interscale–scale architecture, which reflects the spatial organization and lineage output of epidermal stem cell populations in the skin. In homeostatic conditions, Dlx1^+^ slow-cycling stem cells in the interscale region produce a K10^+^ lineage, whereas Slc1a3^+^ fast-cycling stem cells in the scale generate a K31/K36^+^ lineage (Fig. 1A)^16^. Whole-mount immunostaining of epidermal sheets revealed that TPA treatment significantly expanded the size of the K10^+^ interscale region (Fig. 1I, K) while reducing the K36^+^ scale region (Fig. 1J, L), suggesting an alteration in spatial compartments associated with slow- and fast-cycling stem cell populations. Similar changes were observed in imiquimod (IMQ)-treated skin, a model of psoriatic skin disease^53^, suggesting that these changes represent a more common response to inflammatory stress (Fig. S1).

To determine whether this alteration in the epidermal stem cell compartment is permanent or reversible, we allowed mice to recover for 4 weeks after the TPA treatments (Fig. S2A). Epidermal thickness (Fig. S2B, C) and the size and distribution of the interscale and scale regions (Fig. S2D, E) returned to baseline levels, comparable to those of untreated controls. These results indicate that acute inflammatory stress transiently disrupts the spatial organization of epidermal stem cell compartments and that this remodeling is reversible after inflammation resolution.

### Inflammatory stress differentially alters population-specific stem cell behaviors

We performed lineage tracing using compartment-specific CreER drivers to determine whether distinct epidermal stem cell populations differentially respond to inflammatory stress. Dlx1- and Slc1a3-CreER mouse lines were used to label slow- and fast-cycling epidermal stem cell populations, respectively (Fig. 2A; Fig. S3A)^16^. During homeostasis, the Dlx1^+^ population is confined to the K10^+^ interscale lineage, whereas the Slc1a3^+^ population contributes to the K36^+^ scale and interscale line (Fig. 1A). We hypothesized that the reduction in scale size observed under TPA-induced inflammation could result from: (1) a shift in the balance between self-renewal and differentiation of Slc1a3^+^ fast-cycling stem cells toward differentiation, resulting in reduced maintenance of the scale compartment, and/or (2) a bias in differentiation of these stem cells toward the interscale lineage (Fig. S3B).

**Figure 2.**
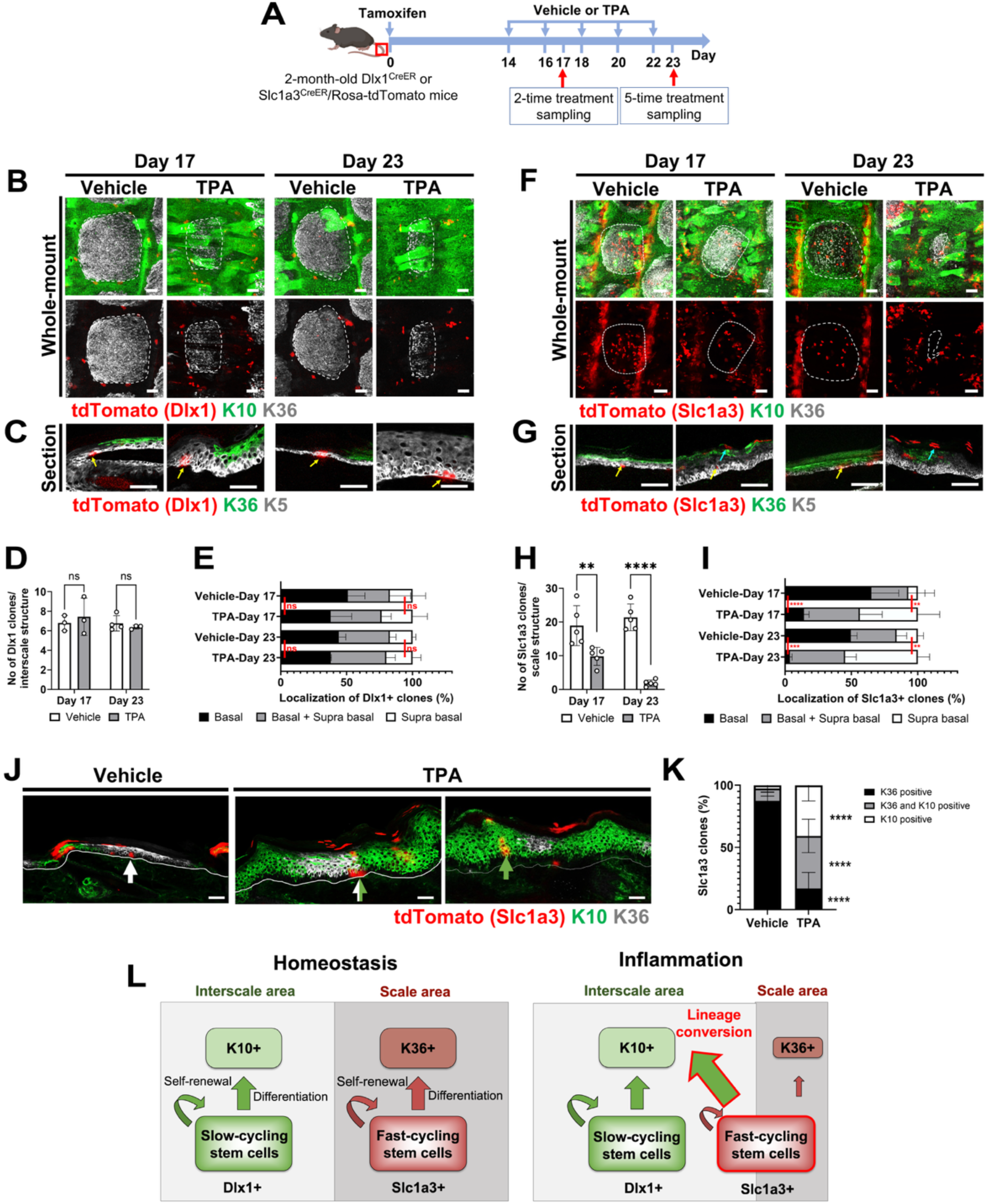
Inflammation differentially alters the lineage dynamics of distinct epidermal stem cell compartments. (**A**) Experimental schedule of lineage tracing. Mice were injected with a single dose of tamoxifen 2 weeks prior to treatment with ethanol (vehicle) or TPA. (**B**) Whole-mount immunostaining of Dlx1-CreER clones with K10 (interscale) and K36 (scale) markers. Scale bars: 100 µm. The white dashed lines indicate the interscale–scale boundary. (**C**) Section immunostaining of Dlx1-CreER^+^ clones with K5 (epidermal basal layer) and K36 (scale) markers. The yellow arrows indicate Tomato^+^ basal clones. Scale bars: 100 µm. (**D**) Number of Dlx1-CreER^+^ clones per one interscale structure. (**E**) Percentage of Dlx1-CreER^+^ clones located in the (1) the basal layer alone (self-renewing clones, black bars); (2) both the basal and suprabasal layers (gray bars); or (3) the suprabasal layer alone (differentiating clones, white bars). (**F**) Whole-mount immunostaining of Slc1a3-CreER^+^ clones with K10 (interscale) and K36 (scale) markers. Scale bars: 100 µm. The white dashed lines indicate the interscale–scale boundary. (**G**) Section immunostaining of Slc1a3-CreER^+^ clones with K5 (epidermal basal layer) and K36 (scale) markers. The yellow and green arrows indicate Tomato^+^ basal and suprabasal clones, respectively. Scale bars: 100 µm. (**H**) Number of Slc1a3-CreER^+^ clones per one interscale structure. (**I**) Percentage of Slc1a3-CreER^+^ clones located in the (1) the basal layer alone (black bars); (2) both the basal and suprabasal layers (gray bars); or (3) the suprabasal layer alone (white bars). (**J, K**) Section immunostaining of Slc1a3-CreER⁺ clones for K10 (interscale) and K36 (scale) in mouse tail skin treated with vehicle (EtOH) or TPA and analyzed on day 23. Scale bars: 100 µm. White arrows indicate Tomato⁺ clones overlapping with K36; green arrows indicate Tomato⁺ clones overlapping with K10; and white–green split arrows denote Tomato⁺ clones overlapping with both K10 and K36. The proportions of Tomato⁺ clones overlapping with K10 (white bar), K36 (black bar), or both markers (gray bar) are quantified. (**L**) Schematic summary of the lineage tracing analysis. Under homeostatic conditions, Dlx1-CreER^+^ slow-cycling epidermal stem cells produce a K10^+^ interscale lineage (green); Slc1a3 produces a K36^+^ scale lineage (red). Under inflammatory conditions, Slc1a3-CreER^+^ clones show enhanced differentiation and a lineage bias toward an interscale-like fate, as evidenced by the ectopic appearance of K10^+^ cells within the scale region. All data are presented as the mean ± SD. Each dot represents an independent biological replicate. Statistical significance was assessed using two-way ANOVA for (D, E, H, I, K). **, *p* < 0.01; ***, *p* < 0.001; ****, *p* < 0.0001; ns, not significant.

We found that Dlx1^+^ clones remained predominantly in the K10^+^ interscale region under both control and inflammatory conditions (Fig. 2B, D). Notably, their abundance, spatial localization, and distribution across basal and suprabasal layers remained unchanged by TPA treatment (Fig. 2C, E). This finding indicates that slow-cycling Dlx1^+^ stem cells maintain self-renewal and differentiation behavior even under inflammatory stress. In contrast, Slc1a3^+^ fast-cycling stem cells showed greater sensitivity to inflammation. TPA treatment led to a progressive reduction in the number of Slc1a3^+^ clones within the scale region (Fig. 2F, H). Although approximately half of Slc1a3^+^ clones remained in the basal layer under control conditions, this proportion declined to ∼15% after two TPA treatments and to <5% after five TPA treatments (Fig. 2G, I). Concomitantly, Slc1a3^+^ clones increased in the suprabasal positions, indicating enhanced differentiation and exit from the basal layer under inflammatory conditions (Fig. 2G, I). Notably, a subset of Slc1a3^+^ clones ectopically expressed K10^+^ interscale markers at the center of the originally scale regions (Fig. 2J, K), suggesting a change in differentiation lineage toward an interscale-like fate. Thus, lineage tracing revealed a differential compartment-specific response: Dlx1^+^ clones remain stable and resilient, whereas Slc1a3^+^ clones undergo inflammation-driven shifts toward differentiation, resulting in reduced basal clone maintenance; concomitantly, a subset of their progeny is biased toward an interscale-like differentiation, together driving tissue-level compartmental remodeling (Fig. 2L).

### Inflammatory stress activates IL-1 signaling and suppresses canonical Wnt signaling in the epidermis

We performed bulk RNA-seq on basal cell populations (α6-integrin^high^/CD34^−^/Sca1^+^) isolated from tail skin after five TPA treatments to uncover the molecular basis underlying the compartment-specific remodeling observed above (Fig. 3A). Principal component analysis (PCA) and hierarchical clustering showed a separation between TPA-treated and control samples (Fig. 3B). Gene ontology (GO) analysis of differentially-expressed genes indicated upregulated expression of inflammation-associated pathways, such as cell cycle regulation, keratinocyte differentiation, cytokine/interleukin signaling, and interleukin-1 beta production regulation (Fig. 3C left; Table S1). In contrast, TPA treatment downregulated the expressions of gene signatures associated with extracellular matrix organization, negative regulation of cell proliferation/differentiation, skin development, and TGF-β and Wnt signaling (Fig. 3C, right; Table S1).

**Figure 3.**
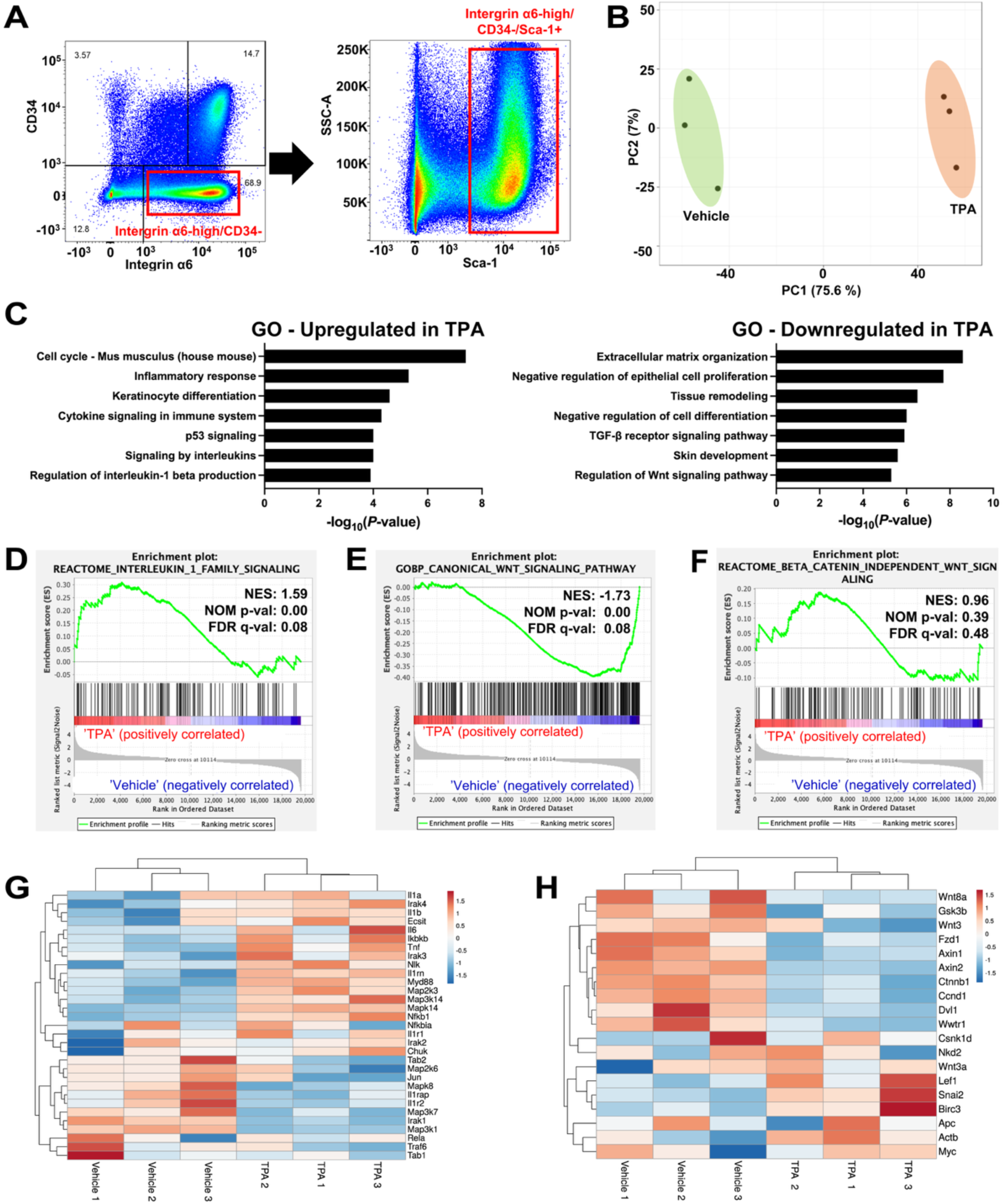
Transcriptomic profiling reveals IL-1 activation and Wnt suppression under inflammatory stress. (**A**) Representative FACS plots. Integrin a6^high^/CD34^−^/Sca1^+^ cells were isolated as epidermal stem cells. (**B**) Transcriptomic profiling of epidermal stem cells isolated from mouse tail skin after five treatments with vehicle (EtOH) or TPA. Principal component analysis of 1.5-fold differentially expressed genes in vehicle (EtOH)-treated and TPA-treated mice. Each dot represents one mouse. (**C**) Gene ontology analysis of 1.5-fold upregulated and downregulated genes in TPA-treated versus vehicle-treated mice. **(D–F)** GSEA showing enrichment of IL-1 family (D), canonical Wnt (E), and non-canonical Wnt (F) signalings in TPA-treated versus vehicle-treated mice. (**G, H**) The heatmap shows changes in IL-1 family (G) and canonical Wnt (H) gene expression.

Among the inflammatory cytokines induced by TPA, IL-1 was the most abundantly induced in the whole tail skin (Fig. 1D), consistent with prior reports implicating IL-1 signaling in TPA-driven epidermal inflammation and hyperplasia^43,54^. Gene set enrichment analysis (GSEA) and heatmap analysis of KEGG-defined pathway genes or GSEA-derived gene sets further indicated the activation of proinflammatory programs, including *Il1a*, *Il1b*, and their receptor *Il1r1*, as well as downstream mediators such as *Myd88*, *Mapk*, *Nfkb1*, *Ikbkb*, *Tnf*, and *Il6* (Fig. 3D, G). Among epidermal stem cell-related pathways, canonical Wnt signaling was suppressed, whereas noncanonical Wnt signatures remained largely unaffected (Fig. 3E, F). The heatmap analysis highlighted reduced expression of multiple canonical Wnt components, including *Wnt3*, *Wnt8a*, *Fzd1*, *Dvl*, *Ctnnb1*, *Axin1/2*, and *Ccnd1* (Fig. 3H). Given that canonical Wnt activity is required for the formation of the fast-cycling scale compartment^15^, its downregulation after TPA treatment potentially contributes to the loss of this population.

Epidermal keratinocytes are the primary producers of the proinflammatory cytokines IL-1α and IL-1β. Furthermore, these cells express high levels of IL-1R1 in the skin^55^. During inflammation, autocrine IL-1 signaling interacts with the EGFR–ERK and integrin pathways, thereby promoting keratinocyte hyperproliferation and amplifying inflammatory responses in the suprabasal layers^43^. In addition, under homeostatic conditions, interfollicular epidermal stem cells self-renew through autocrine Wnt signaling^20^. Therefore, we predicted that IL-1 suppresses canonical Wnt signaling in an epidermis-intrinsic manner, accounting for the suppression of fast-cycling stem cell features. To directly test whether IL-1 signaling is sufficient to suppress canonical Wnt activity in keratinocytes independently of the complex inflammatory milieu induced by TPA, and to assess the conservation of this response in human epidermis, we treated primary human keratinocytes with IL-1β (Fig. 4, Fig. S4) or IL-1α (Fig. S5). Upon IL-1 treatment, proinflammatory cytokines and chemokines (e.g., IL-1α/β, TNF, and CCL20) and IL-1 receptors were robustly induced (Fig. S4; Fig. S5A). Immunoblot analysis revealed strong activation of inflammatory and stress-related signaling pathways, including phosphorylation of NF-κB, JNK, and c-Jun (Fig. 4A, B), whereas AKT and ERK phosphorylation showed no significant changes. Under these conditions, the expression of Wnt target genes, such as *LEF1*, *MYC*, and *CCND1*, was significantly downregulated by IL-1β (Fig. 4C) and IL-1α (Fig. S5B) treatments, consistent with RNA-seq data from TPA-treated mice (Fig. 3H).

**Figure 4.**
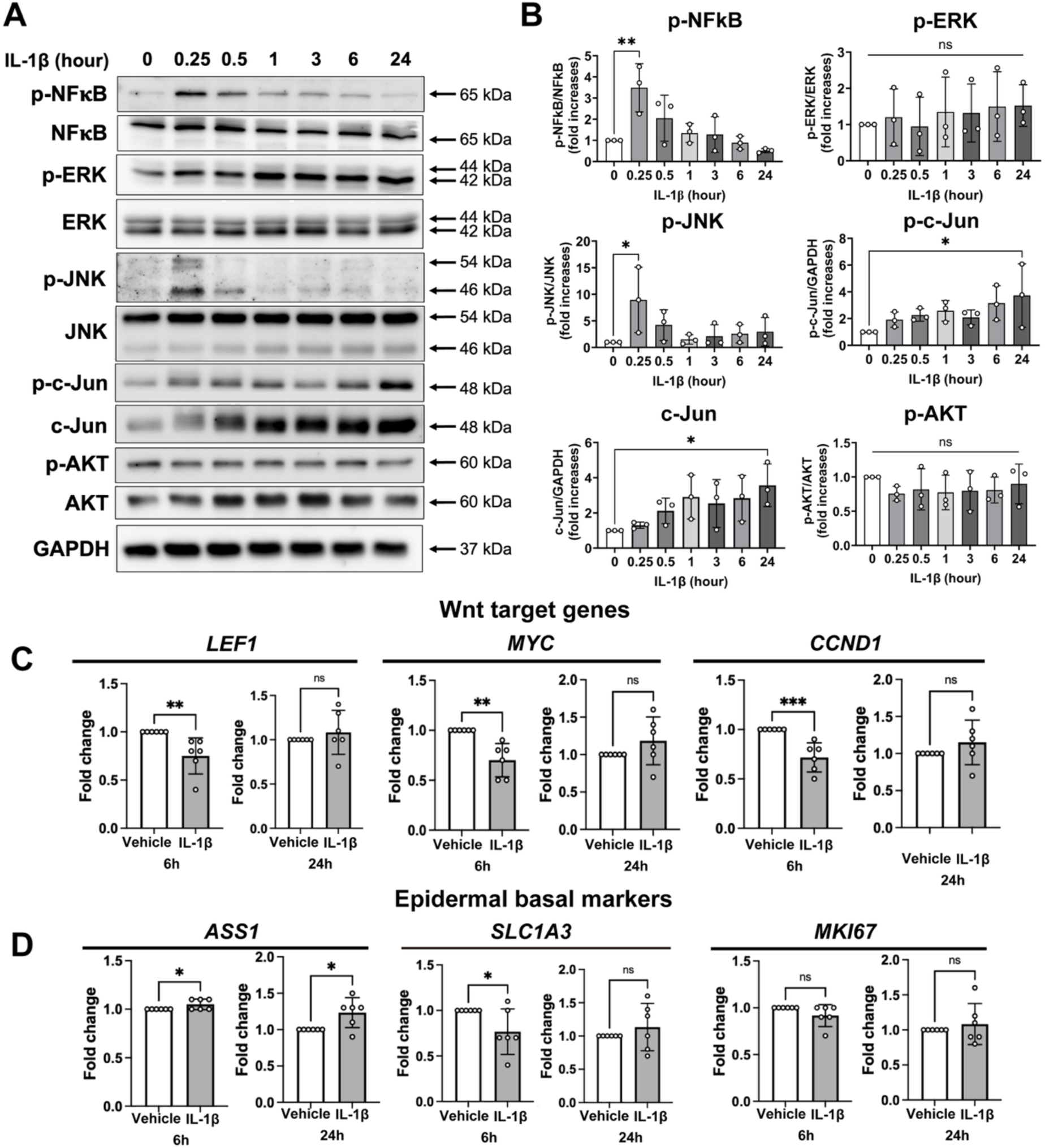
IL-1β suppresses canonical Wnt signaling in primary keratinocytes. **(A)** Neonatal human primary keratinocytes treated with IL-1β over a time course of 24 h were analyzed by western blotting. **(B)** Bar graphs depicting the fold changes of the indicated proteins from (A) after 0.25, 0.5, 1, 3, 6, and 24 hours of IL-1β treatment, compared with unstimulated controls. Each dot represents one sample. **(C, D)** qRT–PCR demonstrating fold changes in Wnt target (C) and epidermal basal marker (D) gene expression after 6 and 24 hours of IL-1β treatment, compared with unstimulated controls. Each dot represents one sample. All data are presented as the mean ± SD. Each dot represents an independent biological replicate. Statistical significance was assessed using one-way ANOVA for (B) and a two-tailed *t* test for (C, D). *, *p* < 0.05; **, *p* < 0.01; ***, *p* < 0.001; ns, not significant.

Human epidermis single-cell RNA-seq studies have demonstrated molecular heterogeneity between epidermal basal cells localized to rete ridges and inter-ridges, which parallels the scale and interscale compartments of murine tail epidermis^30,35,56^. Consistently, IL-1 treatment selectively reduced the expression of the rete ridge-associated marker *SLC1A3* while increasing that of the inter-ridge-associated marker *ASS1*, without affecting *MKI67* levels (Fig. 4D; Fig. S5C). These findings indicate that IL-1 suppresses the rete ridge (the counterpart of scale structures in mice) program while favoring the persistence of inter-ridge compartments. Together, these data suggest that IL-1 suppresses canonical Wnt signaling and disrupts compartment-specific signatures in a keratinocyte-intrinsic manner, consistent with a conserved IL-1–Wnt axis between murine epidermis and human keratinocytes.

### IL-1 signaling governs inflammation-induced remodeling of epidermal stem cell compartments

To test whether *in vivo* IL-1 treatment perturbs epidermal stem cell composition similar to TPA-induced inflammation, we intradermally injected recombinant IL-1α or IL-1β into mouse tail skin (Fig. 5A). Both IL-1α and IL-1β induced an increased expression of proinflammatory cytokines (Fig. S6A, B) and epidermal hyperplasia (Fig. 5B–D). Whole-mount staining revealed a decrease in the K36⁺ scale area and expansion of the K10⁺ interscale region (Fig. 5E–G), mirroring the compartmental changes observed in the TPA model (Fig. 1I–L). These data indicate that IL-1 is sufficient to remodel epidermal stem cell compartments.

**Figure 5.**
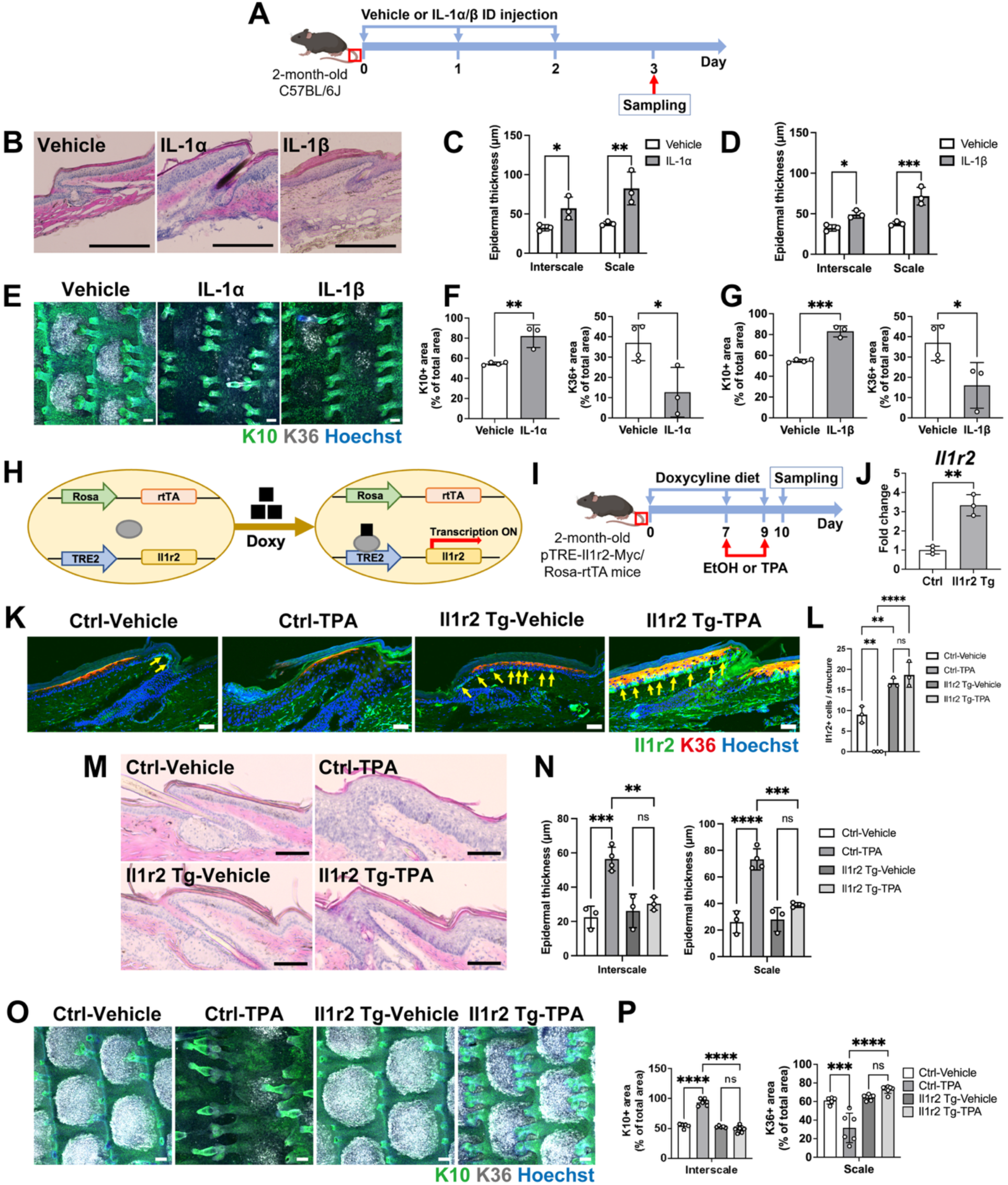
IL-1 signaling is necessary and sufficient for inflammation-induced stem cell compartment remodeling. (**A**) Experimental scheme of IL-1-induced skin inflammation. Mouse tail skin was intradermally (ID) injected with 0.1% BSA (vehicle), IL-1α, or IL-1β once daily for 3 consecutive days. (**B–D**) Hematoxylin and eosin staining of sagittal sections of tail skin (B). Scale bars: 300 µm. Epidermal thickness was measured in the interscale and scale regions (C, D). (**E–G**) Whole-mount staining of K10 (interscale) and K36 (scale), and their quantifications. Scale bars: 100 µm. (**H**) Strategy for overexpressing Il1r2 in pTRE-Il1r2-Myc/Rosa-rtTA transgenic mice (Il1r2 Tg). (**I**) Experimental scheme of Il1r2 overexpression. Il1r2 Tg mice aged 2–3 months were fed doxycycline chow for 7 days before the procedures and maintained on the same diet for the indicated durations. Mouse tail skin was treated with vehicle (EtOH) or TPA on days 7 and 9. (**J**) Il1r2 expression in whole tail skin, measured by qRT–PCR. (**K, L**) Section immunostaining of Il1r2 and K36 (scale) and its quantifications (L). The yellow arrows indicate Il1r2+ cells. Scale bars: 50 µm. The number of Il1r2+ cells per one structure. (**M, N**) Hematoxylin and eosin staining of sagittal sections of tail skin. Scale bars: 100 µm. Epidermal thickness was measured in the interscale and scale regions (N). (**O, P**) Whole-mount staining of K10 (interscale) and K36 (scale), and their quantifications (P). Scale bars: 100 µm. All data are presented as the mean ± SD. Each dot represents an independent biological replicate. Statistical significance was assessed using two-way ANOVA for (C, D), one-way ANOVA for (L, N, P), and a two-tailed *t* test (F, G, J). *, *p* < 0.05; **, *p* < 0.01; ***, *p* < 0.001; ****, *p* < 0.0001; ns, not significant.

IL-1R2 is a decoy receptor that binds IL-1 ligands but does not transduce downstream signals, thereby inhibiting IL-1 signaling^57,58^. RNA-seq analysis revealed that *Il1r2* expression was downregulated following TPA treatment (Fig. 3G), suggesting that inflammatory stress may actively relieve an endogenous suppressor of IL-1 signaling. Thus, we investigated whether blocking IL-1 signaling through enforced IL-1R2 expression could restore the balance between slow- and fast-cycling stem cell compartments, thereby suppressing inflammation-induced remodeling.

To test this hypothesis, we generated a doxycycline-inducible *Il1r2* transgenic mouse line (Fig. 5H, I). qRT–PCR confirmed the increase in *Il1r2* mRNA levels following 1 week of doxycycline administration (Fig. 5J). At the protein level, IL-1R2 was enriched in the slow-cycling interscale compartment, as expected^16^; and the transgene induction resulted in broader IL-1R2 expression extending into the scale region (Fig. 5K, L). Importantly, while TPA treatment reduced IL-1R2 expression in epidermal basal cells of control mice, this reduction was prevented in *Il1r2* transgenic mice (Fig. 5K, L). Functionally, IL-1R2 overexpression attenuated the inflammatory phenotype: levels of IL-1β and IL-6 were reduced (Fig. S6C), epidermal thickening was suppressed (Fig. 5M, N), and the imbalance between scale and interscale compartments was significantly mitigated (Fig. 5O, P). Taken together, these gain-and loss-of-function studies demonstrate that IL-1 is a driver of inflammation-induced remodeling of epidermal stem cell compartments.

### Reactivation of Wnt signaling restores stem cell population balance under inflammatory conditions

To investigate the basis of the preferential impact of inflammatory stress on the fast-cycling stem cell population and its compartmental remodeling, we first examined whether slow- and fast-cycling epidermal stem cells differ in their steady-state signaling activities. Rosa-WntVis reporter mice, which visualize TCF/LEF-dependent canonical Wnt transcriptional activity^59^, were used to analyze the spatial distribution of Wnt-active cells. Reporter expression was highly enriched within the K36⁺ scale compartment; in contrast, overall lower reporter activity was detected in the interscale regions or other skin cell types (Fig. 6A, B). These findings indicate that canonical Wnt signaling is a defining feature of the fast-cycling scale compartment under homeostatic conditions.

**Figure 6.**
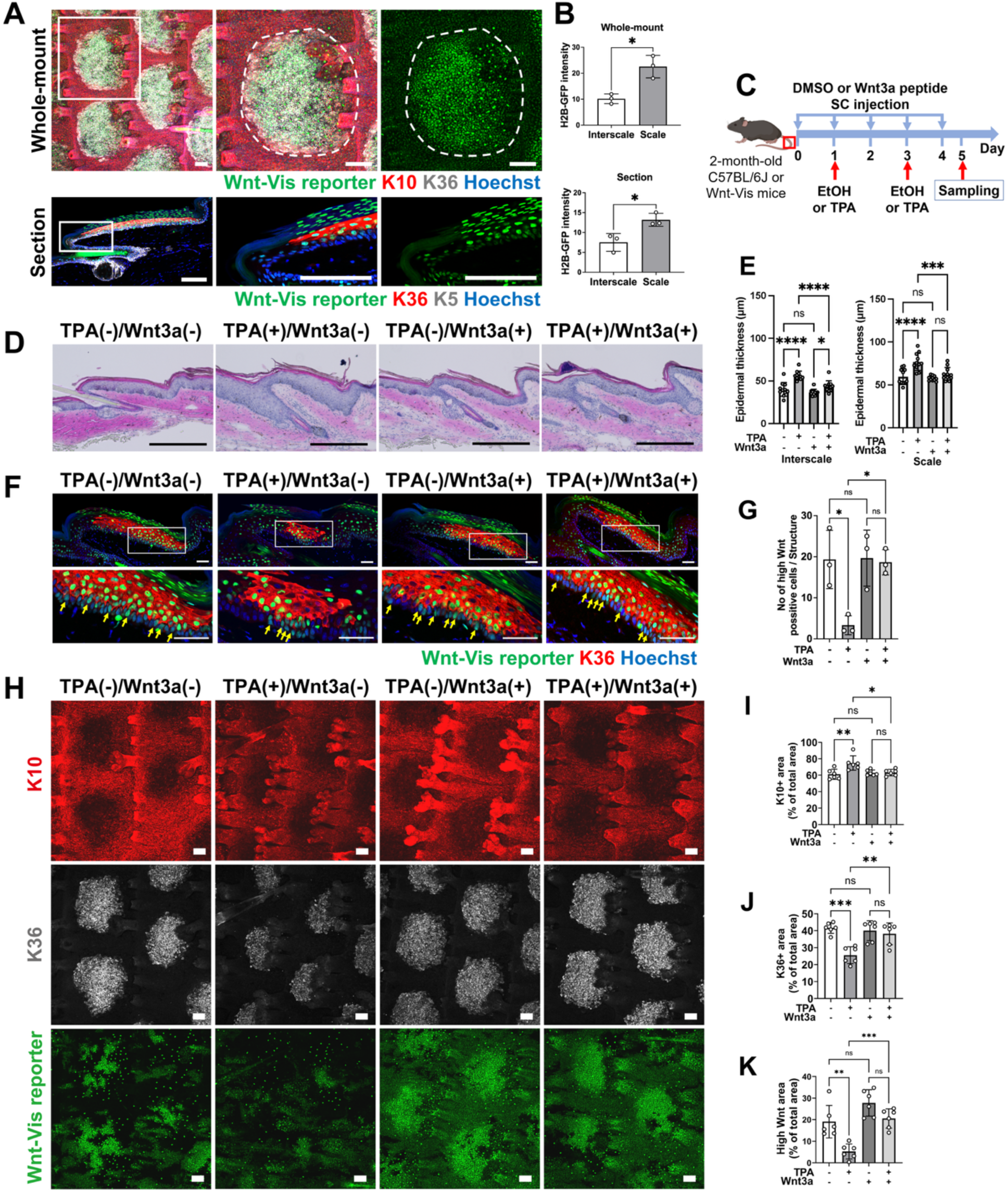
Reactivation of canonical Wnt signaling restores epidermal stem cell balance under inflammation. (**A**) Whole-mount and section immunostaining of Rosa-WntVis reporter mouse tail skin with K10 (interscale), K36 (scale), and K5 (epidermal basal layer) markers. Scale bars: 100 µm. The white dashed lines indicate the interscale–scale boundary. (**B**) Quantification of GFP intensity in interscale and scale regions from whole-mount and sectioned tail skin of Rosa-WntVis reporter mice. (**C**) Experimental scheme. Mouse tail skin was subcutaneously (SC) injected with 1% DMSO (vehicle) or Wnt3a peptide during the topical application with ethanol or TPA. (**D, E**) Hematoxylin and eosin staining of sagittal sections of tail skin (D). Scale bars: 300 µm. Epidermal thickness was measured in the interscale and scale regions (E). (**F, G**) Section immunostaining of Rosa-WntVis reporter with K36 (scale) marker (F). Scale bars: 100 µm. The number of Wnt reporter high cells per one structure (G). (**H–K**) Whole-mount staining of Rosa-WntVis reporter mouse tail skin with K10 (interscale) and K36 (scale) markers (H), with corresponding quantifications shown in (I, J, K). Scale bars: 100 µm. All data are presented as the mean ± SD. Each dot represents an independent biological replicate. Statistical significance was assessed using a two-tailed *t* test for (B) and one-way ANOVA for (E, G, I, J, K). *, *p* < 0.05; **, *p* < 0.01; ***, *p* < 0.001; ****, *p* < 0.0001; ns, not significant.

We next tested whether Wnt signaling reactivation could counteract the loss of the fast-cycling compartment under inflammatory conditions. To this end, we administered a Wnt3a-like peptide (PG-008) subcutaneously during TPA treatment (Fig. 6C). TPA induced epidermal hyperplasia and reduced the K36⁺ scale area, accompanied by suppressed Wnt reporter activity (Fig. 6D–K). Co-treatment with Wnt3a markedly attenuated TPA-induced epidermal hyperplasia (Fig. 6D, E) and prevented the downregulation of Wnt activity, thereby suppressing the remodeling of both compartments, as evidenced by maintenance of the fast-cycling scale compartment (K36⁺) and blunted expansion of the interscale compartment (K10⁺) (Fig. 6F–K). This dual effect suggests that canonical Wnt signaling maintains fast-cycling stem cell behaviors under inflammatory stress, thereby limiting their shifts toward interscale-associated differentiation. Together, these results show that canonical Wnt activity is spatially restricted to the fast-cycling scale compartment and is selectively disrupted under inflammatory stress. Exogenous Wnt activation counteracts this disruption, restoring the scale compartment and rebalancing epidermal stem cell heterogeneity in inflamed skin.

## Discussion

Our study uncovers a previously unrecognized mechanism by which inflammatory cues dynamically reconfigure the composition and spatial organization of epidermal stem cells. Rather than exerting uniform suppression, inflammatory stress elicits population-specific responses, revealing distinct sensitivities and resilience among coexisting stem cell populations. At the cellular level, we demonstrated that inflammatory stress affects fast-cycling epidermal stem cell population by both enhancing their differentiation and biasing their lineage output toward ectopic differentiation into slow-cycling lineages. These observations highlight the dynamic plasticity of this population in adapting its lineage output in response to inflammatory stress, reminiscent of previous studies of lineage plasticity induced by tissue injury^16,60^. Importantly, the change in epidermal stem cell compartments was found to be reversible upon inflammation resolution, suggesting that residual stem cell populations retain the capacity to re-establish their original identities. Although our findings highlight reversible stem cell plasticity during acute inflammation, it remains unknown how these changes may become stabilized or irreversible in chronic inflammation, aging, or cancer.

One key mechanism contributing to this imbalance in stem cell populations is IL-1-mediated suppression of canonical Wnt signaling. Epidermal Wnt/β-catenin activity promotes interfollicular epidermal proliferation^61,62^ and is essential for the formation of fast-cycling stem cell compartments during postnatal skin development^15^. We demonstrated that IL-1 attenuates Wnt activity in keratinocytes without significantly altering AKT phosphorylation (Fig. 4A, B), suggesting that the mechanisms may differ from the AKT-dependent β-catenin degradation described in dermal adipocytes^45^. IL-1 robustly activates NF-κB and JNK/c-Jun (Fig. 4A, B), pathways known to antagonize Wnt/β-catenin signaling, in part by inducing β-catenin inhibitors and sequestering the transcriptional coactivator CBP/p300^63^. In addition, inflammation-associated transcription factors, such as STAT3 and members of the AP-1 family, may further modulate Wnt responsiveness. Future studies dissecting Wnt regulation in a cell type- and context-dependent manner will be required to clarify how these signaling- and transcription-dependent mechanisms control stem cell responses under inflammatory conditions.

The cellular sources and targets of IL-1 and Wnt signals in the epidermis remain an important open question. Epidermal stem cells can secrete Wnt ligands, such as Wnt4 and Wnt10a, to sustain renewal, at least in part, via an autocrine loop^20^. Epidermal keratinocytes^55^, fibroblasts, and multiple immune subsets can produce IL-1α and IL-1β^64^, although their relative contributions during different inflammatory conditions are not well characterized. Epidermal basal cells express IL-1 and Wnt receptors (Fig. 3G, H)^20,55^, indicating that epidermal stem cells integrate these signals directly. Notably, IL-1R2 is selectively expressed in the slow-cycling interscale region but is rapidly downregulated upon TPA treatment (Fig. 5K, L). This observation raises the possibility that IL-1R2 provides an additional, but not exclusive, layer of protection in slow-cycling stem cells, and that their relative resilience may instead reflect intrinsic differences in signaling dependency, such as reduced reliance on Wnt activity. Accordingly, it remains unclear whether the differential sensitivity of fast- and slow-cycling epidermal stem cell populations arises from cell-autonomous differences in signal reception or pathway dependency, or from the environment by surrounding niche cells, including immune and mesenchymal components. Clarifying ligand sources and receptor distributions will be essential to explain these compartment-specific effects.

Beyond these molecular and cellular mechanisms, compartmental specificity also arises from broader tissue organization. Importantly, scale regions critically depend on Wnt signaling, whereas interscale regions are comparatively Wnt-independent^15^. In parallel, melanocytes localize to the scale, whereas Langerhans cells and dendritic epidermal T cells reside in the interscale^65,66^. We previously demonstrated that retinoic acid activation disrupted this spatial organization, inducing ectopic filaggrin expression in scale regions^30^. Aging introduces further remodeling: melanocytes^6^ and dendritic epidermal T cells^67^ decline, and stem cell compartments become impaired^29^. Together, these findings suggest that inflammation-induced IL-1-Wnt crosstalk may intersect with alterations in pigmentation, immunity, skin barrier, and epidermal stem cell dynamics, potentially contributing to physiological and pathological tissue remodeling.

Although the contribution of epidermal stem cell imbalance to tissue pathology remains unclear, recent work has highlighted the emerging roles of epidermal stem cells in tissue renewal, immune surveillance, and the pathogenesis of inflammatory skin disorders^68,69^. Our acute inflammation models demonstrated that IL-1-mediated remodeling of epidermal stem cell compartments is dynamic and reversible; however, chronic or age-associated inflammatory conditions are associated with persistent stem cell dysfunction, as shown by the irreversible impairment of hair follicle stem cells through cytokine imbalance, aberrant JAK–STAT signaling, or the sustained suppression of niche pathways^9,70^. From a disease perspective, these stem cell-level mechanisms may be particularly relevant to chronic inflammatory skin disorders. Transcriptomic analyses have revealed heterogeneous inflammatory responsiveness among basal keratinocyte populations in diseases such as atopic dermatitis and psoriasis^71^, and IL-1β is persistently elevated in psoriatic lesions, where IL-1R signaling correlates with disease progression and therapeutic response^72^. Although blockade of IL-1 family cytokines, including IL-1 and IL-36, provides clinical benefit^73,74^, it remains unresolved whether such interventions normalize epidermal stem cell organization. In parallel, Wnt signaling has emerged as a key regulator of immune balance across tissues, influencing cytokine production and downstream immune responses^75^, in addition to playing a role in stem cell regulation. Taken together, our findings suggest that interventions targeting IL-1 or Wnt signaling may help restore epidermal stem cell equilibrium, raising the prospect of regenerative modulation in chronic inflammatory skin diseases.

### Limitations of the study

Despite the aforementioned findings, several questions remain unresolved. The precise molecular mechanism by which IL-1 suppresses Wnt signaling remains undefined, including effects on ligand availability, β-catenin stability, induction of suppressors, or transcriptional and epigenetic regulation. It is also unclear whether the effects of IL-1-Wnt interactions act directly within epidermal stem cells or indirectly through immune or mesenchymal components. Moreover, the molecular basis of the selective vulnerability of fast-cycling epidermal stem cell populations versus the relative preservation of slow-cycling populations remains unresolved, including whether this heterogeneity reflects differences in IL-1 signaling, Wnt dependency, or an intrinsic transcriptional program. Addressing these questions will require genetic perturbations of IL-1 and Wnt pathways in defined cell populations, integrated with single-cell and spatial multi-omics approaches. Finally, although our acute models reveal reversible changes, the relevance to chronic human disease and aging remains to be established, underscoring the importance of validating these mechanisms in clinical samples and patient-derived datasets.

## Acknowledgments

We would like to thank the Center for Animal Resources and Development (CARD) at Kumamoto University and the Laboratory of Embryonic and Genetic Engineering at Kyushu University for supporting the animal facilities used in this study. We also thank the International Core Facility of Advanced Life Science at Kumamoto University and the Research Promotion Unit at Kyushu University for their core facility support. We thank T. Keida (Kumamoto University) and N. Imai (Kyushu University) for their technical assistance. We thank K. Araki and N. Takeda (Kumamoto University) for their valuable support in generating transgenic mice. This work was supported by the Project for Regenerative, Cellular Medicine and Gene Therapies, AMED (JP25bm1123052) (to A.S. and S.H.), AMED-PRIME, AMED (JP21gm6110016) (to A.S.), Grant-in-Aid for Scientific Research (B) (20H03266, 24K02035) (to A.S.), Grant-in-Aid for Early-Career Scientists (22K15126) (to N.T.K.N.) from the Japan Society for the Promotion of Science (JSPS). This work was also supported by The Takeda Science Foundation (to A.S.), Uehara Memorial Foundation (to A.S.), The Sumitomo Foundation (to A.S.), The Mochida Memorial Foundation for Medical and Pharmaceutical Research (to A.S.), the Lydia O’Leary Memorial Pias Dermatological Foundation (to A.S.), the Inamori Foundation (to A.S.), and The Japan Prize Foundation (to H.T. and A.S.). This work was supported in part by the MEXT Cooperative Research Project Program, the Medical Research Center Initiative for High Depth Omics, and CURE: JPMXP1323015486 for MIB, Kyushu University. Additionally, we acknowledge scholarship support from the Ministry of Education, Culture, Sports, Science, and Technology-Japan (MEXT) (to H.M.P. and T.K.), a SPRING from the Japan Science and Technology Agency (JST), JPMJSP2136 (to I.N.), and the Egypt-Japan Education Partnership (EJEP) Scholarship (to A.M.H.).

## Author contributions

A.S., H.T., and S.H. conceived and designed the study. H.M.P., I.N., N.T.K.N., A.M.H., T.K., and J.A. performed experiments and collected data. H.M.P., I.N., N.T.K.N., A.S., and A.K.Y. conducted formal data analysis and interpretation. H.M.P., I.N., and N.T.K.N. validated the experimental results and curated the datasets. A.S., N.T.K.N., and H.T. developed the methodology and experimental models. A.S., H.T., J.A., and S.F. provided essential resources. A.S. supervised and administered the project. A.S., S.H., N.T.K.N., and H.T. acquired funding. H.M.P., I.N., N.T.K.N., and A.S. contributed to data visualization. H.M.P. and N.T.K.N. wrote the original draft of the manuscript. A.S., I.N., and A.K.Y. reviewed and edited the manuscript. All authors approved the final version of the manuscript.

## Declaration of generative AI and AI-assisted technologies in the writing process

During the preparation of this work, the authors used ChatGPT to assist with clarity and readability. After using this tool, the authors reviewed and edited the content as needed and take full responsibility for the contents of the published article.

## Materials and Methods

### Key resources table

| REAGENT or SOURCE | SOURCE | IDENTIFIER |
| --- | --- | --- |
| **Antibodies** |  |  |
| Ki-67 Monoclonal (SolA15), eBioscience™ | Invitrogen | Cat# 14-5698-82, RRID: AB 10854564 |
| Anti-Cytokeratin 10 [DE-K10] - Cytoskeleton Marker | Abcam | Cat# ab9026, RRID: AB 306950 |
| Cytokeratin 36 Polyclonal | Proteintech | Cat# 14309-1-AP, RRID: AB 2134018 |
| Purified anti-Keratin 5 Polyclonal Chicken | BioLegend | Cat# 905904, RRID: AB 2721743 |
| Phospho-NF-kappaB p65 (Ser536) (93H1) Rabbit Monoclonal | Cell Signaling Technology | Cat# 3033, RRID: AB 331284 |
| NF-kappaB p65 (L8F6) Mouse Monoclonal | Cell Signaling Technology | Cat# 6956, RRID: AB 10828935 |
| Phospho-p44/42 MAPK (Erk1/2) (Thr202/Tyr204) | Cell Signaling Technology | Cat# 9101, RRID: AB 331646 |
| p44/42 MAPK (Erk1/2) | Cell Signaling Technology | Cat# 9102, RRID: AB 330744 |
| Anti-JNK1 + JNK2 + JNK3 (phospho T183+T183+T221) [EPR5693] | Abcam | Cat# ab124956, RRID: AB 10973183 |
| SAPK/JNK | Cell Signaling Technology | Cat# 9252, RRID: AB 2250373 |
| Phospho-c-Jun (Ser73) (D47G9) Rabbit Monoclonal | Cell Signaling Technology | Cat# 3270, RRID: AB 2129575 |
| c-Jun (60A8) Rabbit Monoclonal | Cell Signaling Technology | Cat# 9165, RRID: AB 2130165 |
| Phospho-Akt (Ser473) (736E11) Rabbit Monoclonal | Cell Signaling Technology | Cat# 3787, RRID: AB 331170 |
| Akt | Cell Signaling Technology | Cat# 9272, RRID: AB 329827 |
| GAPDH (14C10) Rabbit Monoclonal | Cell Signaling Technology | Cat# 2118, RRID: AB 561053 |
| BD Horizon™ BV421 Rat Anti-Mouse CD121b (IL1R2) | BD Biosciences | Cat# 562926, RRID: AB 2737896 |
| CD34 Monoclonal Antibody (RAM34), Biotin, eBioscience™ | Thermo Fisher Scientific | Cat# 13-0341-85, RRID: AB 466426 |
| BD Pharmingen™ APC Streptavidin | BD Biosciences | Cat# 554067, RRID: AB 10050396 |
| BD Horizon™ BV421 Rat Anti-Mouse Ly-6A/E | BD Biosciences | Cat# 562729, RRID: AB 2737750 |
| BD Horizon™ BV510 Rat Anti-Human CD49f | BD Biosciences | Cat# 563271, RRID: AB 2738110 |
| BD Horizon™ BV421 Rat IgG2a, κ Isotype Control | BD Biosciences | Cat# 562602, RRID: AB 11153860 |
| BD Horizon™ BV510 Rat IgG2a, κ Isotype Control | BD Biosciences | Cat# 562952, RRID: AB 2869438 |
| Goat anti-Rat IgG (H+L) Cross-Adsorbed Secondary Antibody, Alexa Fluor™ 647 | Invitrogen | Cat# A-21247, RRID: AB_141778 |
| Donkey anti-Mouse IgG (H+L) Highly Cross-Adsorbed Secondary Antibody, Alexa Fluor™ 488 | Invitrogen | Cat# A-21202, RRID: AB_141607 |
| Donkey anti-Rabbit IgG (H+L) Highly Cross-Adsorbed Secondary Antibody, Alexa Fluor™ 647 | Invitrogen | Cat# A-31573, RRID: AB_2536183 |
| Goat anti-Rat IgG (H+L) Cross-Adsorbed Secondary Antibody, Alexa Fluor™ 555 | Invitrogen | Cat# A-21434, RRID: AB_2535855 |
| Donkey anti-Rabbit IgG (H+L) Highly Cross-Adsorbed Secondary Antibody, Alexa Fluor™ 555 | Invitrogen | Cat# A-31572, RRID: AB_162543 |
| Goat anti-Chicken IgY (H+L) Secondary Antibody, Alexa Fluor™ 647 | Invitrogen | Cat# A-21449, RRID: AB_2535866 |
| Donkey anti-Rat IgG (H+L) Highly Cross-Adsorbed Secondary Antibody, Alexa Fluor™ 488 | Invitrogen | Cat# A-21208, RRID: AB_2535794 |
| Donkey anti-Mouse IgG (H+L) Highly Cross-Adsorbed Secondary Antibody, Alexa Fluor™ 555 | Invitrogen | Cat# A-31570, RRID: AB_2536180 |
| Goat Anti-Rabbit IgG (H + L)-HRP Conjugate | Bio-Rad | Cat# 1706515, RRID: AB_11125142 |
| Goat Anti-Mouse IgG (H + L)-HRP Conjugate | Bio-Rad | Cat# 1706516, RRID: AB_11125547 |
| **Chemicals, peptides, and recombinant proteins** |  |  |
| Ethanol | FUJIFILM Wako | 056-03341 |
| Mayer's Hematoxylin Solution | FUJIFILM Wako | 131-09665 |
| Eosin Y | FUJIFILM Wako | 058-00062 |
| Acetic Acid | FUJIFILM Wako | 017-00256 |
| Xylene | FUJIFILM Wako | 244-00081 |
| FastStart Essential DNA Green Master | Roche | 6402712001 |
| RNAprotect Tissue Reagent | Qiagen | 1018087 |
| Sodium Azide | FUJIFILM Wako | 197-11091 |
| Ethylenediamine-N,N,N',N'-tetraacetic Acid Disodium Salt Dihydrate (2NA) | FUJIFILM Wako | 345-01865 |
| 10xD-PBS(-) | FUJIFILM Wako | 048-29805 |
| PBS, pH 7.4 | Thermo Fisher Scientific | 10010023 |
| Paraformaldehyde | FUJIFILM Wako | 162-16065 |
| Bovine Serum Albumin | Sigma-Aldrich | A4503 |
| Normal Donkey Serum | Sigma-Aldrich | D9663 |
| Normal Goat Serum | Sigma-Aldrich | G9023 |
| Triton™ X-100 | Nacalai Tesque, INC. | 35501-15 |
| TWEEN® 20 | Sigma-Aldrich | P1379 |
| Hoechst 33342 | Sigma-Aldrich | B2261 |
| p-Phenylenediamine | FUJIFILM Wako | 164-01532 |
| Glycerol | FUJIFILM Wako | 072-00626 |
| Phorbol 12-myristate 13-acetate | Sigma-Aldrich | P8139 |
| IMQ | Mochida Pharmaceutical | 224130002 |
| Vaseline | Unilever | N/A |
| Entellan™ new rapid mounting medium for microscopy | Sigma-Aldrich | 107961 |
| Tamoxifen | Sigma-Aldrich | T5648 |
| Glycine | FUJIFILM Wako | 077-00735 |
| Gelatin from cold water fish skin | Sigma-Aldrich | G7765 |
| Coin oil | Sigma-Aldrich | C8267 |
| Acetone | FUJIFILM Wako | 013-00356 |
| Trypsin | MP Biomedicals | 210314080 |
| 45w/v% D(+)-Glucose solution | FUJIFILM Wako | 079-005511 |
| DMEM, high glucose | Thermo Fisher Scientific | 11965092 |
| Cytiva HyClone™ US Bovine Calf Serum | GE healthcare Life Science | SH3007303 |
| PE Annexin V Apoptosis Detection Kit with 7-AAD | BioLegend | 640934 |
| TRIzol™ LS Reagent | Invitrogen | 10296-028 |
| Tissue-Tek® O.C.T. Compound | Sakura Finetek | 4583 |
| BD Horizon™ Brilliant Stain Buffer | BD Biosciences | 563794 |
| Cnt-PR | CELLnTEC | CEL-1100-00 |
| Accutase Cell Detachment Solution | Innovative Cell Technologies, Inc. | AT104-500 |
| Accumax Cell Detachment Solution | Innovative Cell Technologies, Inc. | AM105-500 |
| Human Collagen Type IV | Sigma-Aldrich | C5533 |
| 0.4w/v% Trypan Blue Solution | FUJIFILM Wako | 207-17081 |
| RIPA Buffer | Sigma-Aldrich | R0278 |
| Protease Inhibitor Cocktail | Sigma-Aldrich | P8340 |
| Phosphatase Inhibitor Cocktail SolutionI (×100) | FUJIFILM Wako | 167-24381 |
| 2-Mercaptoethanol, 99% | FUJIFILM Wako | 131-14572 |
| 2-Amino-2-hydroxymethyl-1,3-propanediol | FUJIFILM Wako | 015-20093 |
| Hydrochloric Acid | FUJIFILM Wako | 080-01066 |
| Sodium Hydroxide | FUJIFILM Wako | 198-13765 |
| Sodium Chloride | Nacalai Tesque, INC. | 31333-45 |
| Ammonium Peroxodisulfate | Nacalai Tesque, INC. | 02627-21 |
| 40(w/v)%-Acrylamide/Bis Mixed Solution(29:1) | Nacalai Tesque, INC. | 06119-45 |
| N,N,N',N'-Tetramethylethylenediamine | FUJIFILM Wako | 205-06313 |
| Methanol | Sigma-Aldrich | 34860 |
| ECL™ Advance Blocking Reagent | Sigma-Aldrich | GERPN418 |
| ECL™ Prime Western Blotting System | Sigma-Aldrich | GERPN2232 |
| Sodium Dodecyl Sulfate | FUJIFILM Wako | 194-13985 |
| Mouse IL-1 alpha Recombinant Protein, PeproTech® | Thermo Fisher Scientific | 211-11A |
| Mouse IL-1 beta Recombinant Protein, PeproTech® | Thermo Fisher Scientific | 211-11B |
| Human IL-1 alpha Recombinant Protein, PeproTech® | Thermo Fisher Scientific | 200-01A |
| Human IL-1 beta Recombinant Protein, PeproTech® | Thermo Fisher Scientific | 200-01B |
| Doxycycline food | Oriental Kobo | Custom order |
| PG-008 Wnt3a alternative peptide (β-catenin pathway agonist) | PeptiGrowth | PG-008 |
| PageRuler™ Prestained Protein Ladder, 10 to 180 kDa | Thermo Fisher Scientific | 26616 |
| Bromophenol Blue | FUJIFILM Wako | 021-02911 |
| Domicum | Maruishi Pharma | 211—762100 |
| Vetorphale | Meiji Seika Pharma | 4987970000000 |
| Saline | Otsuka Pharmaceutical Factory, Inc. | 873311 |
| Dormitor | Nippon Zenyaku Kogyo | 14111 |
| **Critical commercial assays** |  |  |
| RNeasy Mini Kit | Qiagen | 74104 |
| RNeasy Micro Kit | Qiagen | 74004 |
| iScript™ Reverse Transcription Supermix | Bio-Rad | 1708841 |
| M.O.M.® (Mouse on Mouse) Immunodetection Kit | Vector Laboratories, Inc. | BMK-2202 |
| Pierce™ BCA Protein Assay Kits | Thermo Fisher Scientific | 23227 |
| Immunoreaction Enhancer Solution Can Get Signal™ | Toyobo | NKB-101 |
| **Deposited data** |  |  |
| Bulk RNA-seq data | N/A | GEO: GSE312274 |
| **Experimental models: Cell lines** |  |  |
| Neonatal human primary keratinocytes | BIOPREDIC International | KER112006 |
| **Experimental models: Organisms/strains** |  |  |
| Mouse: C57BL/6J | Charles River Laboratories/Japan SLC |  |
| Mouse: Dlx1-CreER (C57BL/6J) | Jackson Laboratory | Stock no. 014551 |
| Mouse: Slc1a3-CreER (C57BL/6J) | Jackson Laboratory | Stock no. 012586 |
| Mouse: Rosa-tdTomato reporter | Jackson Laboratory | Stock no. 007905 |
| Mouse: Rosa-WntVis reporter | RIKEN BRC | RBRC10162 |
| Mouse: pTRE-Il1r2-Myc | Sada Laboratory | N/A |
| Mouse: Rosa-rtTA | Jackson Laboratory | Stock no. 005670 |
| **Vector and Oligonucleotides** |  |  |
| pTRE vector | Clontech | 631008 |
| Primers | Sigma-Aldrich | Custom oligos |
| **Software and Algorithms** |  |  |
| Fuji (Image J) | Fuji (Image J) | https://fiji.sc |
| FlowJo v10 | BD Biosciences | https://flowjo.com/flowjo10/download |
| Adobe photoshop | Adode software | https://www.adobe.com/products/photoshop.html |
| Adobe illustrator | Adode software | https://www.adobe.com/products/illustrator.html |
| GraphPad Prism 10 | Dotmatics | www.graphpad.com |
| Biorender | Biorender | https://www.biorender.com |
| GSEA | BROAD institute, UC San Diego | https://www.gsea-msigdb.org/gsea/index.jsp |
| ClustVis | ClustVis | https://biit.cs.ut.ee/clustvis/ |
| Metascape | Zhou et al. Nature Commun. 2019 10(1):1523 | https://metascape.org/gp/index.html#/main/step1 |
| CLC Genomics Workbench Premium | QIAGEN | https://digitalinsights.qiagen.com/products-overview/discovery-insights-portfolio/analysis-and-visualization/qiagen-clc-genomics-workbench/ |

## Methods

### Mice

Wild-type (C57BL/6J) mice were purchased from Charles River Laboratories or Japan SLC, Inc. For the lineage-tracing experiment, Dlx1-CreER (C57BL/6J) (The Jackson Laboratory, no. 014551)^76^, Slc1a3-CreER (C57BL/6J) (The Jackson Laboratory, no. 012586)^77^ mice were crossed with Rosa-tdTomato reporter mice (C57BL/6J) (The Jackson Laboratory, no. 007905)^78^. Rosa-WntVis reporter mice were used to assess canonical Wnt activity^59^.

Experiments were conducted using mice of both sexes, aged 2–3 months unless otherwise specified, with group allocation performed randomly. The mice were housed in specific-pathogen-free (SPF) facilities with a 12-hour light/dark cycle and *ad libitum* access to food and water. The mice were provided with routine care, including weekly cage changes.

### Ethical statement

All animal experiments were approved by the Animal Care and Use Committee of Kumamoto University and the Animal Experiment Committee of Kyushu University, and were conducted in accordance with institutional guidelines for animal experimentation.

### Human primary keratinocytes

Neonatal human primary keratinocytes (KER112006) were obtained from BIOPREDIC International (Saint-Grégoire, France), where they were collected with written informed consent and provided to investigators in anonymized form in accordance with applicable ethical standards. Keratinocytes with passage number less than 10 were cultured in CnT-PR medium (CELLnTEC) at 37°C in 5% CO2 on 100-mm dishes pre-coated with collagen IV (Sigma-Aldrich) until they reached 70–80% confluency.

### Generation of pTRE-Il1r2 transgenic mice

pTRE-Il1r2 transgenic mice were generated using a tetracycline-inducible (Tet-On) system. The Il1r2 coding sequence with a C-terminal Myc tag was PCR-amplified and cloned into the multicloning site of the pTRE vector (Clontech) using BamHI and HindIII restriction sites. The expression cassette was excised from the plasmid by XhoI and AseI digestion and purified for pronuclear microinjection. The microinjection into C57BL/6J zygotes was performed at the Center for Animal Resources and Development (CARD) of Kumamoto University. Genomic integration was confirmed by PCR with pTRE-Il1r2 transgene-specific primers (F 5′-TCGCCTGGAGACGCCAT-3′, R 5′-ACTGACAGTTGTCTCCATGGA-3′). pTRE-Il1r2 mice were crossed with Rosa-rtTA mice (JAX no. 005670). Double-transgenic mice (aged 2–3 months) received doxycycline chow (1 g/kg, Oriental Kobo Inc.) for 7 days prior to downstream procedures and continuously thereafter for the indicated durations. A total of nine founder lines were obtained; among them, lines #4 and #9 showed stable germline transmission across generations and reproducible Il1r2 induction and phenotypes upon doxycycline administration.

### 12-O-tetradecanoylphorbol-13-acetate (TPA) treatment and recovery experiments

To induce local inflammation, TPA (32 nmol; Sigma-Aldrich) diluted in 99.5% ethanol (100 µL) was applied topically to the mouse tail skin. Control mice received ethanol alone. Applications were performed either two or five times at 2-day intervals, and skin was harvested 1 day after the last treatment. For recovery experiments, mouse tail skin was topically treated with 100 µl of TPA (32 nmol, Sigma-Aldrich) or ethanol for control mice, five times at 2-day intervals. Skin tissues were collected for analysis after a 4-week recovery period.

For pTRE-Il1r2-Myc/Rosa-rtTA mice, TPA (8 nmol) diluted in ethanol was applied by pipette at 80 µl to ∼4 cm² of tail skin. For Rosa-WntVis reporter mice, 40 µl of the TPA solution (4 nmol) was applied to ∼2 cm² of tail skin. Control mice received equivalent volumes of ethanol (80 µl or 40 µl, respectively). Skin applications were performed twice every 2 days, and skin tissues were harvested 1 day later.

### Imiquimod (IMQ) treatment

To establish the alternative acute inflammation model, 100 µl of acetone was applied topically, followed by 5% IMQ cream (Beselna, Mochida Pharmaceutical) to approximately 4 cm² of tail skin (62.5 mg/day) once daily for 7 consecutive days. For controls, acetone was applied followed by vehicle (Vaseline) cream. Skin was collected 1 day after the last application.

### Induction of inflammation via the application of IL-1α and IL-1β to mouse tail skin

The intradermal IL-1 injection procedure was performed on mouse tail skin as previously described^54,79^, with some modifications. Recombinant murine IL-1α (Thermo Fisher Scientific) and IL-1β (Thermo Fisher Scientific) were reconstituted in sterile PBS with 0.1% bovine serum albumin (BSA). IL-1α or IL-1β (125 ng/25 µl per site) was intradermally injected into 4 sites within a ∼2 cm² tail skin once daily for 3 days using 30-gauge needles. Controls mice received an equivalent volume of PBS with 0.1% BSA. Skin tissues were harvested 1 day later.

### Subcutaneous injection of Wnt3a-like peptide

The subcutaneous injection into the mouse tail skin was performed as previously described, with minor modifications^80^. A Wnt3a-like peptide (Peptidream/PeptiGrowth PG-008; 20 ng in 20 µl per site in PBS/0.1% BSA/1% DMSO) or vehicle was subcutaneously injected into 4 sites within a ∼2 cm² tail skin region once daily for 5 days. On days 2 and 4, 40 µl of TPA (4 nmol) or ethanol was applied 6 hours after Wnt3a/vehicle treatment. Skin was collected on day 6 (1 day after the final injection).

### Tamoxifen and doxycycline administration

Tamoxifen (Sigma–Aldrich) was injected intraperitoneally at a single dose of 50 μg/g body weight for Slc1a3-CreER and Dlx1-CreER mice. For the overexpression of Il1r2, pTRE-Il1r2/Rosa-rtTA mice were fed doxycycline chow (1 g/kg, Oriental Kobo Inc.) beginning 1 week before TPA treatment. This feed was maintained throughout the experiment.

### Whole-mount immunostaining of tail epidermal sheets

Skin tissue was cut into 5 mm × 5 mm cubes and incubated in 20 mM EDTA on a shaker at 37°C for 2 hours, after which the epidermis and dermis were separated using tweezers. The separated epidermal sheet was fixed overnight in 4% PFA at 4°C. After washing with 1× PBS, the epidermal sheet was stored in PBS with 0.2% sodium azide at 4°C. After blocking with a blocking buffer (1% BSA, 2.5% NDS, 2.5% NGS, 0.8% Triton in PBS) for 3 hours, the primary antibody was incubated overnight. The primary antibodies used were as follows: anti-rat Ki-67 (1:100, Invitrogen), anti-mouse Keratin 10 (1:500, Abcam), and anti-rabbit Keratin 36 (1:300, Proteintech). After primary antibody incubation, the samples were incubated with 0.2% Tween in PBS for 1 hour at room temperature, then washed four times. Secondary antibody incubation was performed overnight at 4°C. Secondary antibodies (Alexa 488/647/555, Invitrogen) were used at a dilution of 1:500. The samples were incubated with 0.2% Tween in PBS for 1 hour at room temperature and washed four times, and then stained with Hoechst 33342 (Sigma-Aldrich, 1:1000) in 1× PBS for 1 hour at room temperature. To remove excess Hoechst, the sections were washed with 1× PBS for 15 minutes and mounted with an anti-fade mounting medium. When using mouse antibodies, the MOM kit (Vector Laboratories) was used according to the manufacturer’s instructions^81^. Images were acquired on a Nikon A1 and a Zeiss LSM 900 using NIS-Elements AR and ZEN 2010 software, respectively. All whole-mount images are presented as maximum-intensity projections of Z-stacks from the basal side.

### Immunostaining of tail skin tissue sections

Tail skin tissue was embedded and frozen in OCT compound (Sakura Finetech Japan Co., Ltd.), then cut into 10 µm-thick tissue sections. The sections were fixed in 4% PFA at room temperature for 10 minutes. The sections were then washed three times with 1× PBS, three times with 20 mM Glycine/PBS, and once with 0.1% Triton/PBS for 5 minutes each. Blocking was performed with blocking solution (1% BSA, 2.5% NDS, 2.5% NGS, 2% Gelatin, 0.1% Triton in PBS) for 1 hour at room temperature. The sections were then stained with primary antibodies overnight at 4 °C. The primary antibodies used were as follows: anti-rabbit Keratin36 (1:500, Proteintech), anti-chicken Keratin5 (1:500, BioLegend), and Anti-Mouse CD121b (IL-1R2) (BD Biosciences). Secondary antibodies (Alexa Fluor 488/555/647, Invitrogen; 1:500) were applied for 1 hour at room temperature. Nuclei were counterstained with Hoechst 33342 (Sigma-Aldrich, 1:1000). Slides were mounted in anti-fade medium and imaged as described above.

### Hematoxylin and eosin staining

Tail skin was directly embedded in OCT compound (Tissue-Tek, Sakura). Ten-micrometer sections were fixed in 4% paraformaldehyde at room temperature for 10 minutes. The sections were then stained with hematoxylin (Wako) for 3 minutes and eosin Y (Wako) for 15 seconds, followed by dehydration and mounting in Entellan solution (Merck Millipore).

### Quantitative RT-qPCR of whole mouse skin

Approximately 5 mg of frozen tail skin tissue, stabilized in RNAprotect Tissue Reagent (Qiagen) overnight at 4°C, was pulverized under liquid nitrogen using a precooled mortar and pestle. The powdered tissue was immediately resuspended in 600 µL RLT buffer supplemented with 1% β-mercaptoethanol and homogenized by repeated passage through a 26-gauge needle. Total RNA was extracted from frozen mouse tail skin using an RNeasy Mini Kit (Qiagen) according to the manufacturer’s instructions. RNA concentration and purity were assessed by measuring 1 µl of RNA solution with a NanoDrop spectrophotometer (Thermo Fisher Scientific). cDNA was synthesized using the iScript cDNA Synthesis Kit (Bio-Rad). qRT–PCR was performed on a LightCycler 96 System (Roche) using FastStart Essential DNA Green Master (Roche) with 5 ng of cDNA per reaction. Target gene expression was normalized to the Actb housekeeping gene, and relative expression levels were calculated with the control group set to 1. The primer sequences used for qRT–PCR are as follows: Actb: forward (F) 5′-GATCAGCAAGCAGGAGTACGA-3′, reverse (R) 5′-AAAACGCAGCTCAGTAACAGTC-3′; Il1a: F 5′-CTGGAAGAGACCATCCAACCC-3′, R 5′-ACTTCCTGTTGCAGGTCATTT-3′; Il1b: F 5′-CCCTGCAGCTGGAGAGTGTGGA-3′, R 5′-TGTGCTCTGCTTGTGAGGTGCTG-3′; Tnfa: F 5′-GCCCACGTCGTAGCAAACCAC-3′, R 5′-GCAGGGCTCTTGACGCAG-3′; Il6: F 5′-CCAGTTGCCTTCTTGGGACTG-3′, R 5′-GCCATTGCACAACTCTTTTCTC-3′; and Il1r2: F 5′-CCCCTGGAGACAATACCAGC-3′, R 5′-TTAGCCAACCACCACACAATG-3′.

### FACS isolation

Tail skin from wild-type (C57BL/6J) mice treated five times with ethanol or TPA (32 nmol) was used for FACS isolation and subsequent RNA-seq analysis. Subcutaneous and fat tissues were removed from the tail skin, which was then incubated overnight at 4°C in 0.25% trypsin/EDTA and, the following day, incubated at 37°C for 30 minutes. Single-cell suspensions were prepared by gently scraping the epidermis and filtering through 70 μm and 40 μm filters. Cells were stained on ice for 30 minutes with the following antibodies: CD34-biotin (1:50, eBioscience, 13-0341), Streptavidin-APC (1:100, BD Biosciences, 554067), α6-integrin-BUV395 (1:100, BD Biosciences, custom order), and Sca1-BV421 (1:100, BD Biosciences, 562729). Dead cells were excluded using 7-AAD (BioLegend). Cells were isolated using a FACS Aria flow cytometer (BD Biosciences). Single-color controls were used for gate setting and compensation. The data were analyzed with FlowJo software (BD Biosciences).

### Bulk RNA-sequencing and analysis

Cells isolated by FACS were directly sorted into TRIzol LS (Ambion, 10296028) and sent to Tsukuba i-Laboratory LLP, University of Tsukuba, for further analysis. RNA integrity was assessed using an Agilent 2100 Bioanalyzer. RNA-seq libraries were prepared with the SMARTer® Stranded Total RNA-Seq Kit v2 – Pico Input Mammalian (Takara) and sequenced on the NextSeq500 platform (Illumina). Data were analyzed using CLC Genomics Workbench 11 (Qiagen). Genes exhibiting a fold change ≥1.5 were selected for downstream analyses. PCA and heatmaps were conducted using online tools including Morpheus (Morpheus, https://software.broadinstitute.org/morpheus) and Clustvis^82^. GSEA and GO analyses were conducted using GSEA software^83,84^ and Metascape^85^, respectively.

### Keratinocyte treatments, qRT–PCR, and western blotting

Human primary keratinocytes were passaged using 2 ml Accutase (ICT) and 1 ml Accumax (ICT) per 100 mm dish for 15–20 minutes at room temperature, then seeded onto collagen IV-coated 12-well plates at 5 × 10⁴ cells/well. The cells were then cultured in CnT-PR medium at 37°C in 5% CO_2_ for 24 hours, then switched to medium with IL-1α/β (10 ng/mL, Thermo Fisher Scientific) and cultured for 0.25, 0.5, 1, 3, 6, and 24 hours.

Total RNA from cultured human keratinocytes was isolated with the RNeasy Micro Kit (QIAGEN). cDNA synthesis and qRT–PCR were performed as described above for mouse tail skin. The primer sequences were as follows: *SLC1A3* forward (F) 5’-AGCAGGGAGTCCGTAAACGC-3’ and reverse (R) 5’-TGGTCGGAGGGTAAATCCAAGG-3’; *ASS1* F 5’-TCTACAACCGGTTCAAGGGC-3’ and R 5’-TCCAGGATTCCAGCCTCGTA-3’; *LEF1* F 5’-CCCGTGAAGAGCAGGCTAAA-3’ and R 5’-GCAGCTGTCATTCTTGGACCT-3’; *MYC* F 5’-GCTGCTTAGACGCTGGATTT-3’ and R 5’-CACCGAGTCGTAGTCGAGGT-3’; *CCND1* F 5’-AATGACCCCGCACGATTTC-3’ and R 5’-TCAGGTTCAGGCCTTGCAC-3’; *MKI67* F 5’-ATCCAGCTTCCTGTTGTGTCA-3’ and R 5’-GCTGGCTCCTGTTCACGTAT-3’; *ACTB* F 5’-AGAGATGGCCACGGCTGCTT-3’ and R 5’-ATTTGCGGTGGACGATGGAG-3’; *IL1A* F 5’-GCCATCGCCAATGACTCAGA-3’ and R 5’-ATGTAATGCAGCAGCCGTGA-3’; *IL1B* F5’-CCTGAGCTCGCCAGTGAAAT-3’ and R 5’-GGTGGTCGGAGATTCGTAGC-3’; TNF F 5’-GACAAGCCTGTAGCCCATG-3’ and R 5’-GGACCTGGGAGTAGATGAGG-3’; *CCL20* F 5’-GCGAATCAGAAGCAGCAAGC-3’ and R 5’-GATTTGCGCACACAGACAACT-3’; *IL1R1* F5’-GCTCATCGTGATGAATGTGGC-3’ and R 5’-ACTGGCCGGTGACATTACAG-3’; and *IL1R2* F 5’-TAGGACGGTCCCAGGAGAAG-3’ and R 5’-AACGGCAGGAAAGCATCTGT-3’.

For western blotting, cells were washed with PBS and lysed in 1× RIPA buffer (Sigma-Aldrich) supplemented with phosphatase (FUJIFILM Wako) and protease (Sigma-Aldrich) inhibitor cocktails. Protein concentrations were measured using a BCA Protein Assay Kit (Thermo Fisher Scientific), and samples were mixed with 4× SDS sample buffer containing 10% 2-mercaptoethanol (FUJIFILM Wako). The samples were then heated at 95 °C for 10 minutes and cooled to room temperature. Equal protein concentrations (5 µg/lane) were subsequently separated by SDS–PAGE. The proteins were then transferred to methanol-activated PVDF membranes (Merck Millipore).

The membranes were blocked with 5% ECL Advance Blocking Reagent (Sigma-Aldrich) containing 0.1% Tween-20 (TBS-T) for 1 hour, washed, and incubated overnight at 4 °C with primary antibodies diluted in immunoreaction enhancer solution 1 (Toyobo). The primary antibodies used were as follows: anti-rabbit phospho-NF-kappaB p65 (Ser536) (93H1) (1:1000, Cell Signaling), anti-mouse NF-kappaB p65 (L8F6) (1:1000, Cell Signaling), anti-rabbit phospho-p44/42 MAPK (Erk1/2) (1:4000, Cell Signaling), anti-rabbit p44/42 MAPK (Erk1/2) (1:4000, Cell Signaling), anti-rabbit JNK1 + JNK2 + JNK3 (phospho T183+T183+T221) [EPR5693] (1:1000, Abcam), anti-rabbit SAPK/JNK (1:1000, Cell Signaling), anti-rabbit phospho-c-Jun (Ser73) (D47G9) (1:1000, Cell Signaling), anti-rabbit c-Jun (60A8) (1:1000, Cell Signaling), anti-rabbit phospho-Akt (Ser473) (736E11) (1:2000, Cell Signaling), anti-rabbit Akt (1:2000, Cell Signaling), and anti-rabbit GAPDH (14C10) (1:3000, Cell Signaling). After washing, the membranes were incubated for 1 hour with HRP-conjugated secondary antibodies (Bio-Rad) diluted in immunoreaction enhancer solution 2 (Toyobo). Subsequently, signals were developed using the ECL Prime Western Blotting System (Sigma–Aldrich) and imaged with an ImageQuant LAS 4000 mini (GE Healthcare). Band intensities were quantified using ImageJ (Fuji) and normalized to the control.

### Quantification and statistical analysis

#### Imaging data

In H&E-stained tail sections, an epidermal unit was defined as the interfollicular epidermis (IFE) region encompassing one scale and one interscale structure, typically located between two hair follicles (HFs). Epidermal thickness was measured from six epidermal units per mouse^86^, with scale IFE thickness assessed at the center of an epidermal unit and interscale IFE thickness measured adjacent to the HF. K10/K36-positive areas, the number of tdTomato-positive cell clones, and clonal areas were manually quantified from 4–8 images representing 16–32 interscale or scale IFE structures per mouse, using projected Z-stack images analyzed with ImageJ (NIH). GFP signal intensity in the tail skin of Rosa-WntVis reporter mice was quantified according to the mean gray value. The average GFP intensity of basal cells within the K36-positive region was used as the threshold to define high Wnt-positive cells in tissue sections.

#### Statistical analyses

Quantifications were independently conducted on at least three mice. Analyses were performed unblinded due to practical constraints (limited personnel). Data are presented as mean ± standard deviation (SD), and each dot in the graphs represents an independent biological replicate. A two-tailed unpaired Student’s *t* test (two groups) or one or two-way ANOVA (≥3 groups) was used as appropriate (GraphPad Prism v10.x). Definition of significance: ∗*p* < 0.05, ∗∗ *p* < 0.01, ∗∗∗ *p* < 0.001, ∗∗∗∗ *p* < 0.0001, ns: not significant. Outlier exclusion (ROUT, Q = 1%) was applied only when pre-defined criteria were met and was performed identically across experimental groups.

---

**Figure S1.**
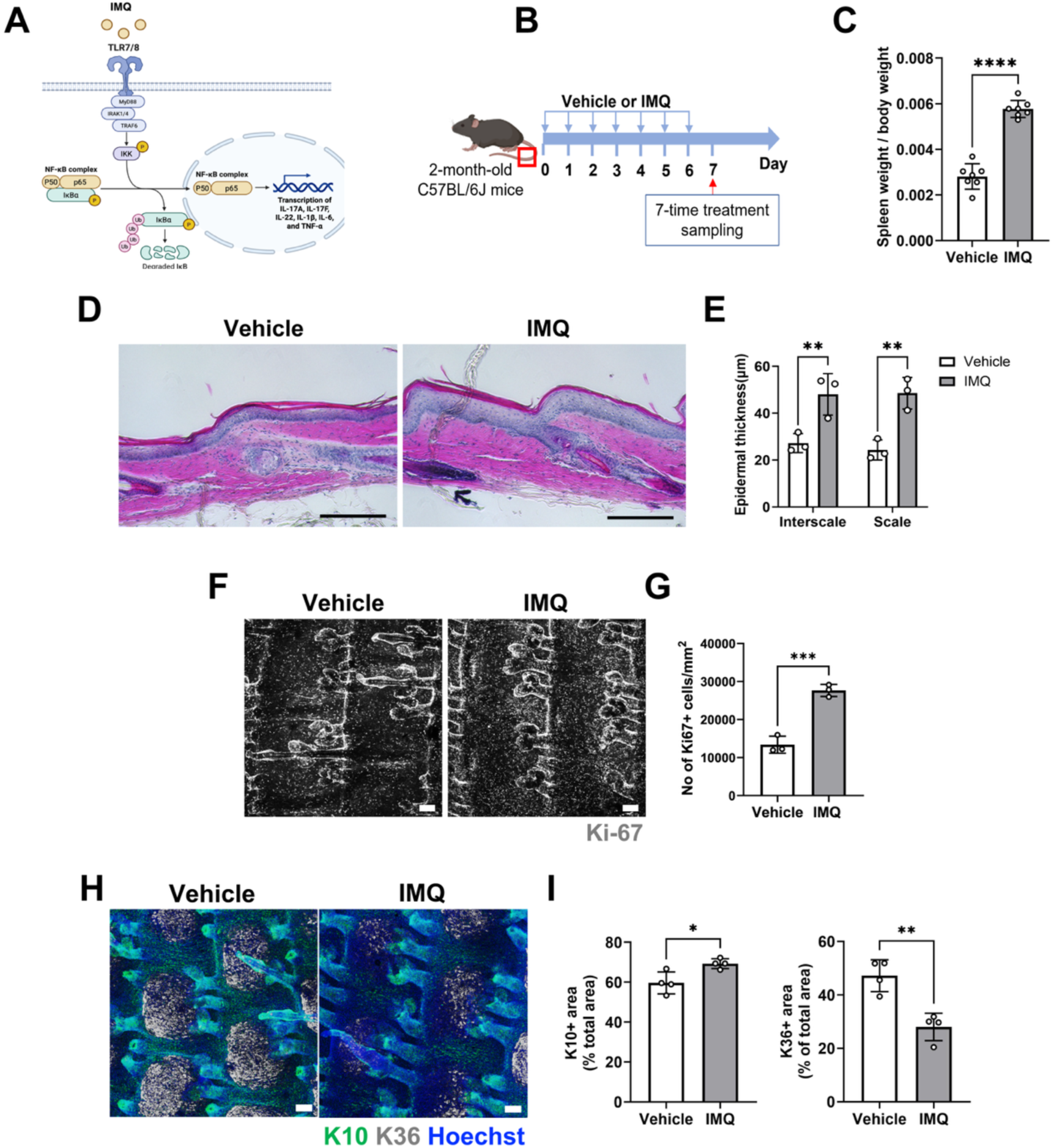
Imiquimod-induced inflammation remodels epidermal compartments. (**A**) Molecular pathway of Imiquimod (IMQ)-induced skin inflammation. (**B**) Experimental scheme of IMQ-induced skin inflammation. Mouse tail skin was treated topically with vaseline (vehicle) or IMQ for 7 consecutive days and dissected one day after the last treatment. (**C**) Spleen weight/body weight. (**D, E**) Hematoxylin and eosin staining of sagittal sections of tail skin. Scale bars: 300 µm. Epidermal thickness was measured in the interscale and scale regions (E). (**F, G**) Whole-mount immunostaining of Ki-67 (F) and its quantification (G). Scale bars: 100 µm. The data are presented as the mean ± SD. Each dot represents one mouse. (**H, I**) Whole-mount staining of K10 (interscale) and K36 (scale), and their quantifications (I). Scale bars: 100 µm. All data are presented as the mean ± SD. Each dot represents an independent biological replicate. Statistical significance was assessed using a two-tailed *t* test for (C, G, I) and two-way ANOVA for (E). *, *p* < 0.05; **, *p* < 0.01; ***, *p* < 0.001; ****, *p* < 0.0001.

**Figure S2.**
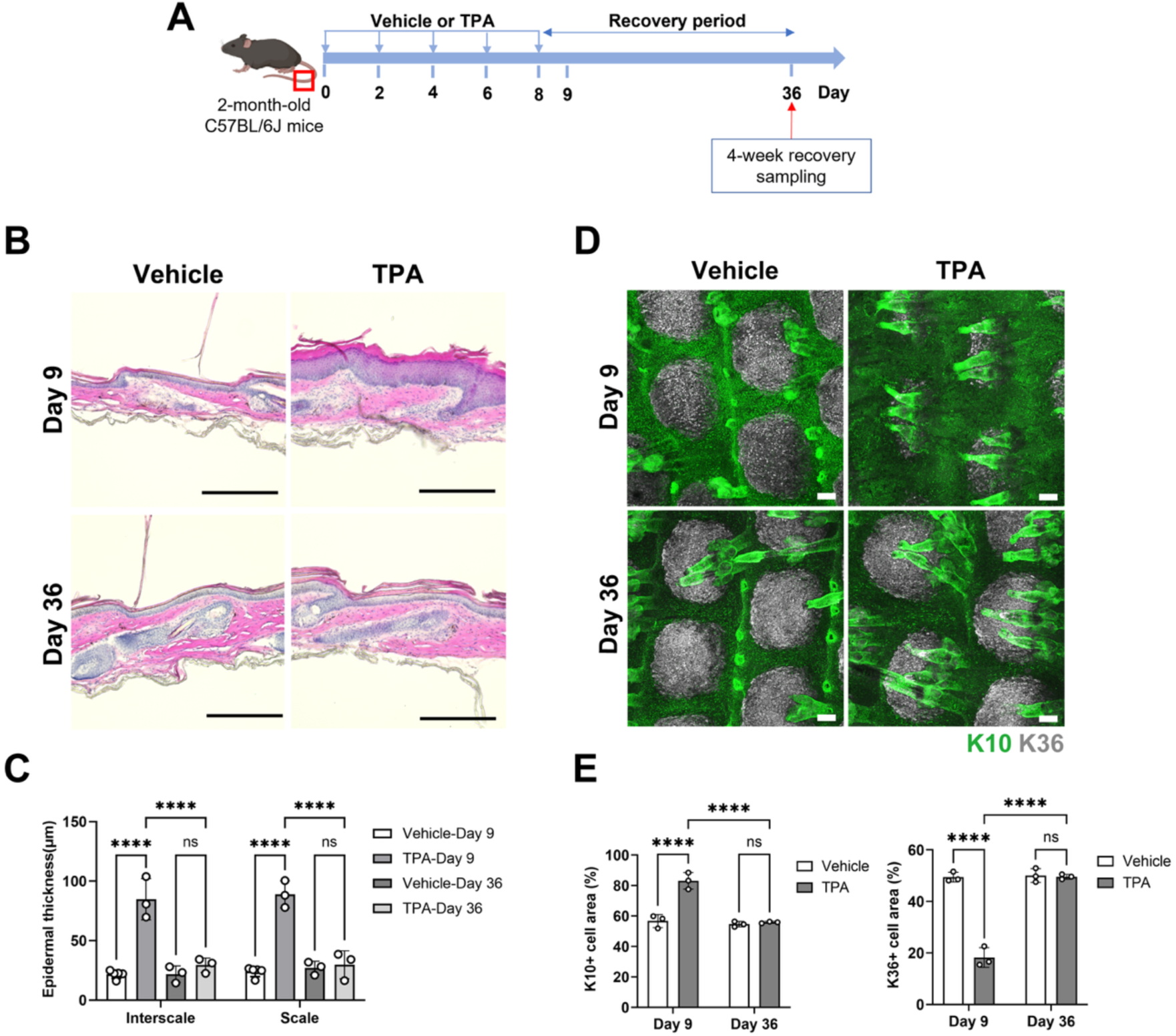
Epidermal stem cell compartment remodeling is reversible after TPA withdrawal. (**A**) Experimental scheme of recovery experiments. (**B, C**) Hematoxylin and eosin staining of sagittal sections of tail skin. Scale bars: 300 µm. Epidermal thickness (white arrows) was measured in the interscale and scale regions (C). The data are presented as the mean ± SD. Each dot represents one mouse. (**D–E**) Whole-mount staining of K10 (interscale) and K36 (scale), and their quantifications (E). Scale bars: 100 µm. All data are presented as the mean ± SD. Each dot represents an independent biological replicate. Statistical significance was assessed using a two-way ANOVA for (C, E). ****, *p* < 0.0001; ns, not significant.

**Figure S3.**
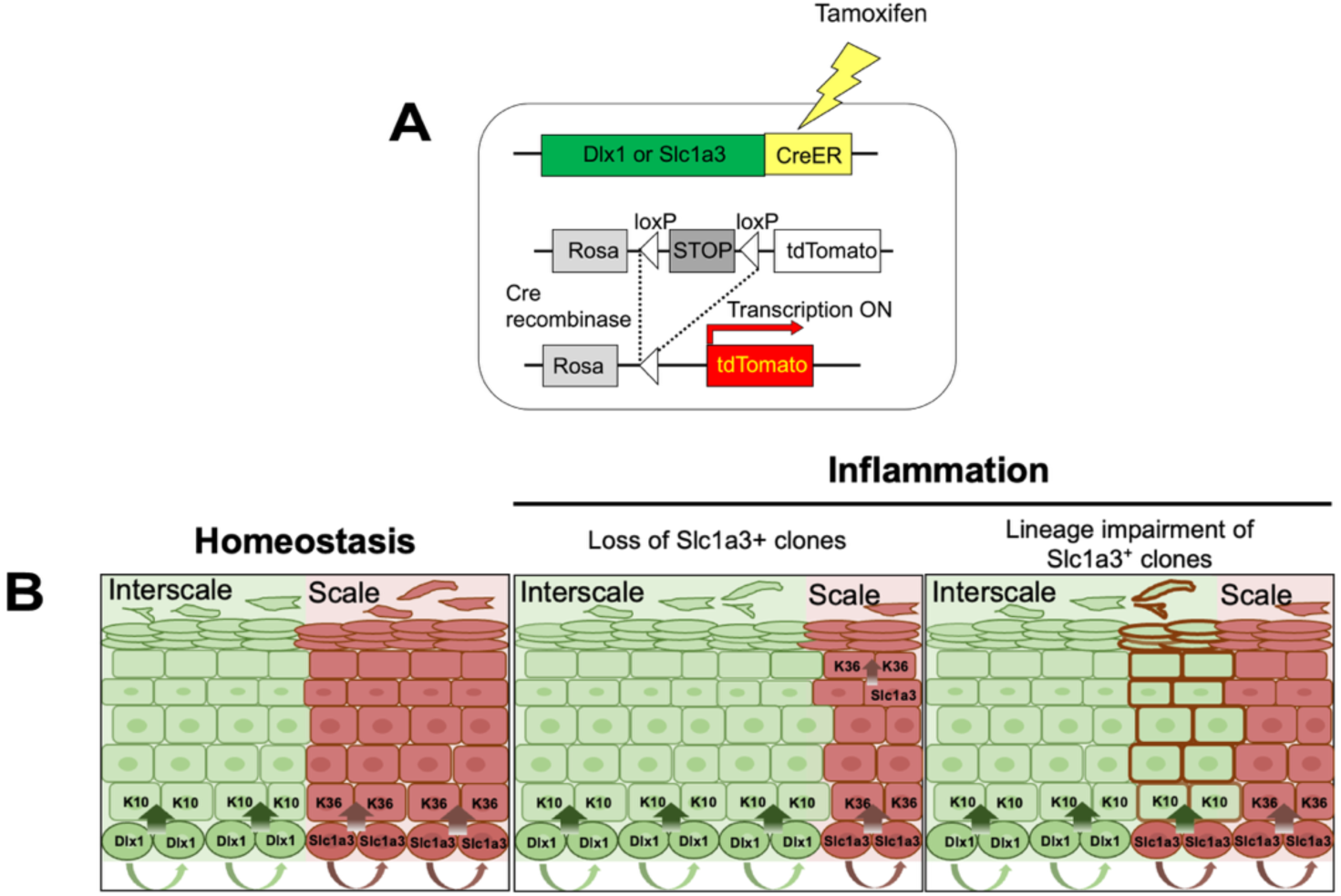
Schematic representation of the lineage tracing strategy. (**A**) Diagram illustrating the tamoxifen-inducible CreER system. (**B**) Potential behaviors of Dlx1^+^ slow-cycling and Slc1a3^+^ fast-cycling epidermal stem cell populations under homeostatic or TPA-treated conditions.

**Figure S4.**
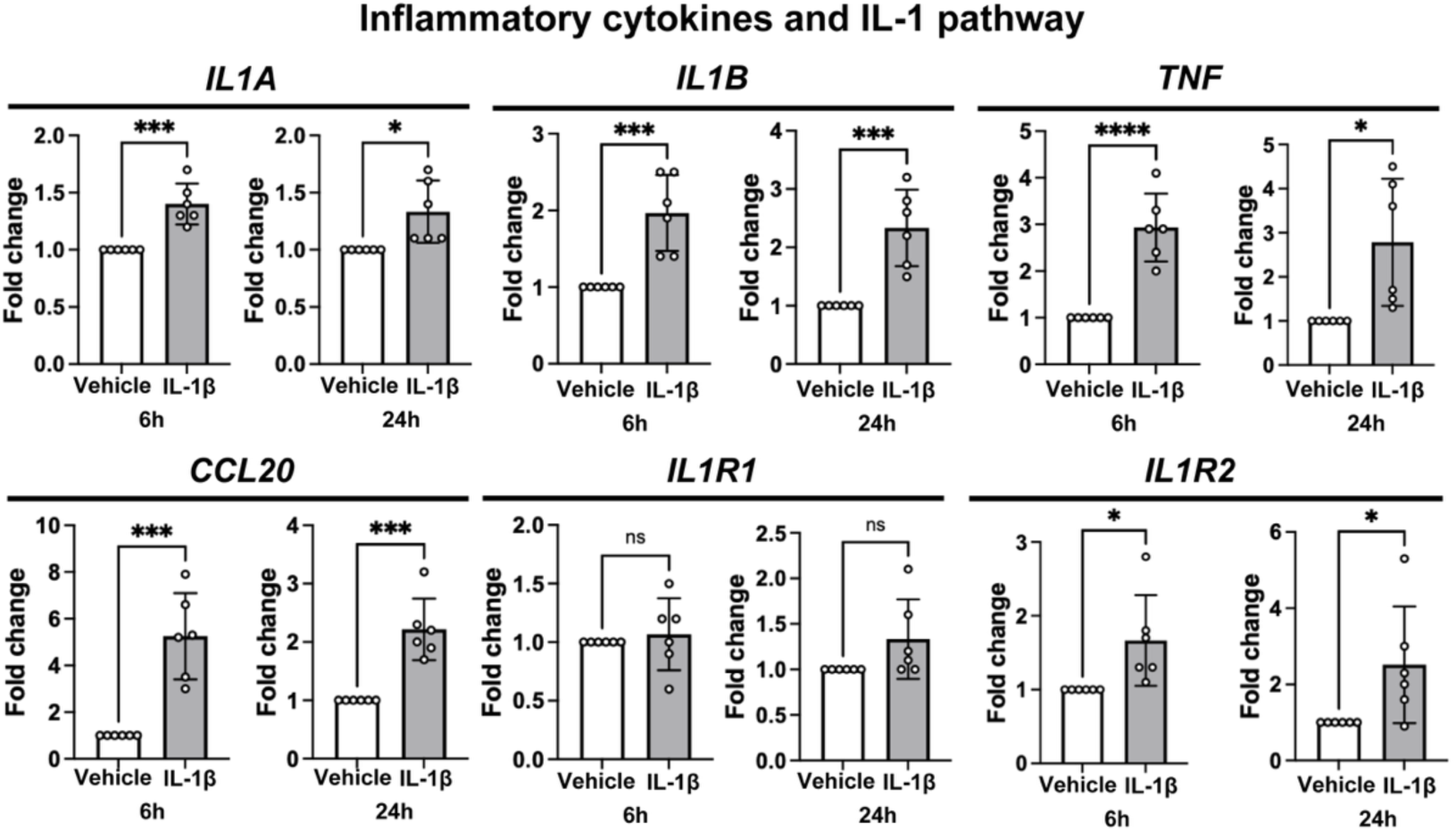
IL-1β-induced inflammation in neonatal human primary keratinocytes. qRT–PCR showing fold changes in inflammatory cytokines and IL-1 pathway-related genes after 6 or 24 hours of IL-1β treatment compared with unstimulated controls. Data are presented as the mean ± SD. Each dot represents an independent biological replicate. Statistical significance was assessed using a two-tailed *t* test: *, *p* < 0.05; ***, *p* < 0.001; ****, *p* < 0.0001; ns, not significant.

**Figure S5.**
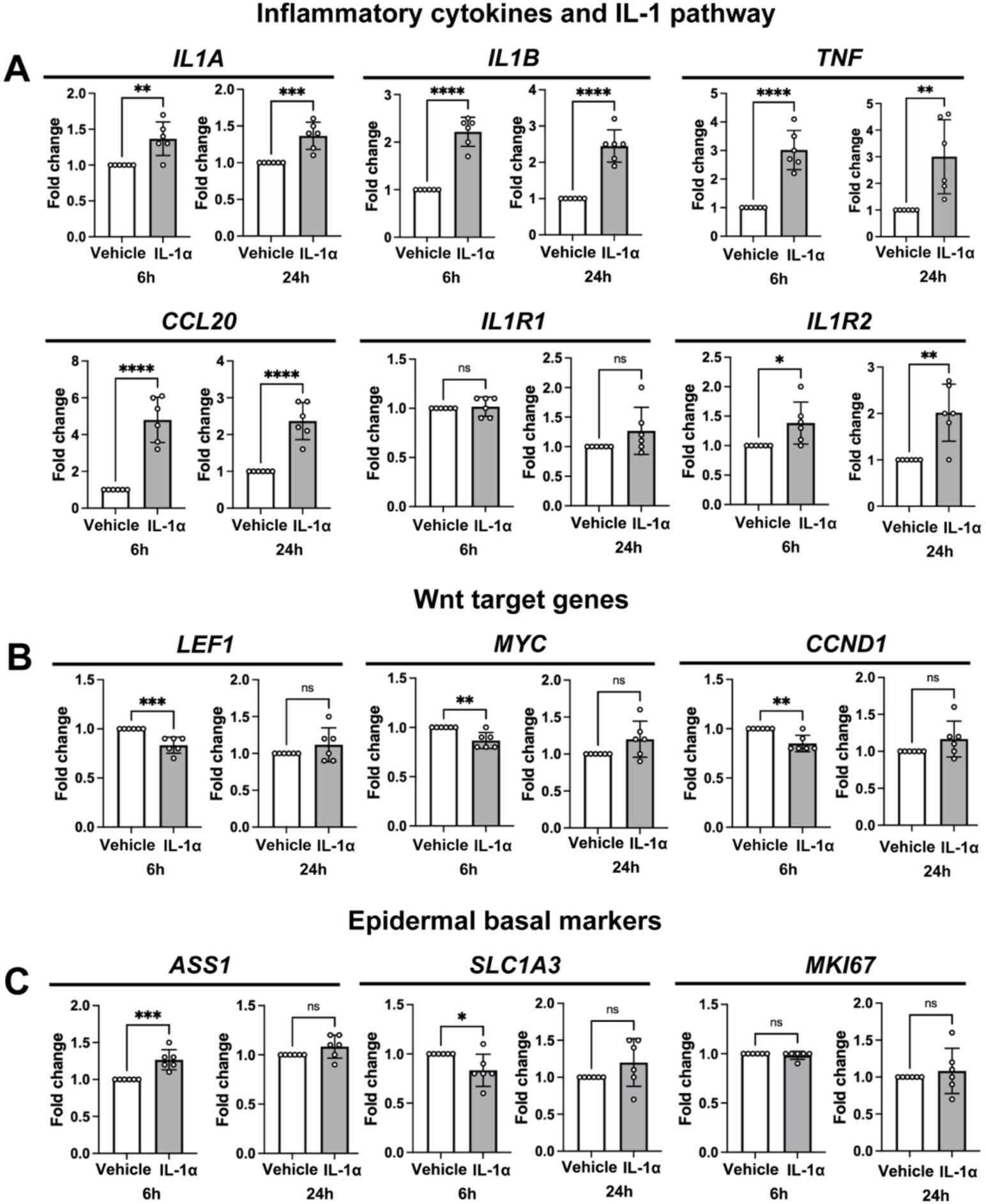
IL-1α suppresses canonical Wnt activity in primary keratinocytes. (**A–C**) qRT–PCR showing fold changes in inflammatory cytokines and IL-1 pathway genes (A), Wnt target genes (B), and epidermal basal markers (C) after 6 or 24 hours of IL-1α treatment compared with unstimulated controls. Data are presented as the mean ± SD. Each dot represents an independent biological replicate. Statistical significance was assessed using a two-tailed *t* test: *, *p* < 0.05; **, *p* < 0.01; ***, *p* < 0.001; ****, *p* < 0.0001; ns, not significant.

**Figure S6.**
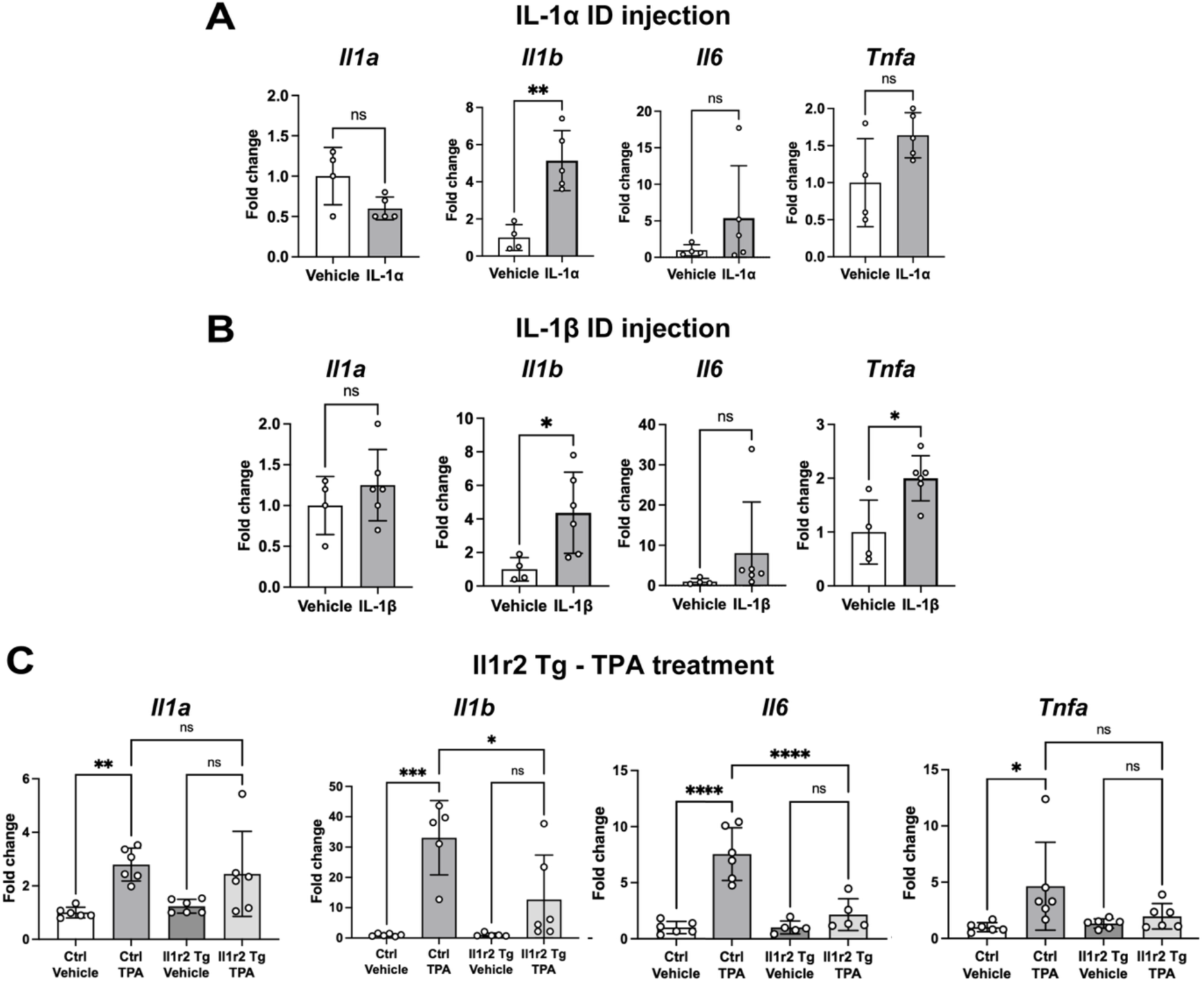
Cytokine expression changes upon IL-1 gain- and loss-of-function. (**A, B**) Expression of inflammatory cytokines in the whole tail skin of WT mice injected with IL-1α (A), or IL-1β (B) once daily for 3 consecutive days, measured by qRT-qPCR. (**C**) Expression of inflammatory cytokines in the whole tail skin of pTRE-Il1r2-Myc/Rosa-rtTA (Il1r2 Tg) or control (Ctrl) mice treated twice with ETOH or TPA, measured by qRT–PCR. All data are presented as the mean ± SD. Each dot represents an independent biological replicate. Statistical significance was assessed using a two-tailed *t* test for (A, B) and one-way ANOVA for (C). *, *p* < 0.05; **, *p* < 0.01; ***, *p* < 0.001; ****, *p* < 0.0001; ns, not significant.

